# Tuft cells restrain pancreatic tumorigenesis through paracrine eicosanoid signaling

**DOI:** 10.1101/2019.12.19.882985

**Authors:** Kathleen E. DelGiorno, Chi-Yeh Chung, H. Carlo Mauer, Sammy Weiser Novak, Rajshekhar R. Giraddi, Dezhen Wang, Razia F. Naeem, Linjing Fang, Leonardo R. Andrade, Nikki K. Lytle, Wahida H. Ali, Crystal Tsui, Vikas B. Gubbala, Maya Ridinger-Saison, Makoto Ohmoto, Carolyn O’Connor, Galina A. Erikson, Maxim Nikolaievich Shokhirev, Yoshihiro Urade, Ichiro Matsumoto, Vera Vavinskaya, Pankaj K. Singh, Uri Manor, Kenneth P. Olive, Geoffrey M. Wahl

**Affiliations:** Gene Expression Laboratory, Salk Institute for Biological Studies, La Jolla, CA, 92037; Department of Medicine, Herbert Irving Comprehensive Cancer Center, Columbia University Irving Medical Center, New York, NY, 10032; Waitt Advanced Biophotonics Center, Salk Insitute for Biological Studies, La Jolla, CA, 92037; Eppley Institute for Research in Cancer, University of Nebraska Medical Center, Omaha, NE, 68198; Monell Chemical Senses Center, Philadelphia, PA, 19104; Flow Cytometry Core, Salk Insitute for Biological Studies, La Jolla, CA, 92037; Razavi Newman Integrative Genomics and Bioinformatics Core, Salk Institute for Biological Studies, La Jolla, CA, 92037; Isotope Science Center, The University of Tokyo, Bunkyo-ku, Tokyo 113-0032, Japan; Department of Pathology, University of California San Diego, San Diego, CA, 92103; Klinik und Poliklinik für Innere Medizin II, Klinikum rechts der Isar, Technical University, 81675 Munich, Germany

## Abstract

Despite numerous advances in our understanding of pancreatic ductal adenocarcinoma (PDA) genetics and biology, this disease is expected to become the second leading cause of cancer-related U.S. deaths within the next few years. Incomplete understanding of how it arises precludes development of early detection and interception strategies to improve therapeutic outcomes. Acinar to ductal metaplasia involving genesis of tuft cells is one early step in PDA formation, but their functional significance has remained obscure due to their rarity and a lack of methods and relevant animal models for their molecular and functional analysis. Here, we show that deletion of tuft cell master regulator Pou2f3 eliminates pancreatic tuft cells and increases fibrosis, alters immune cell activation, and accelerates disease progression. We demonstrate that tuft cell expression of the prostaglandin D_2_ synthase Hpgds restrains pancreatic disease progression in early stages by inhibiting stromal activation. Analyses of human data sets are consistent with mouse studies. We propose that tuft cells and, by inference, the associated metaplastic lesions, play a protective role early in pancreatic tumorigenesis.

**Significance:** We find that tuft cell formation in response to oncogenic *Kras* is protective and restrains tumorigenesis through local production of anti-inflammatory substances, including paracrine prostaglandin D_2_ signaling to the stroma. Our findings establish tuft cells as a metaplasia-induced tumor suppressive cell type.

## Introduction

Chemical or mechanical injury of the pancreas results in a cell state-switching event termed acinar to ductal metaplasia (ADM) where digestive enzyme producing acinar cells dedifferentiate into proliferative ductal-like cells to restore pancreas homeostasis. This process is perturbed during neoplastic progression, where, in mouse models, initiating *Kras* mutations prevent tissue healing and, instead, ADM is thought to progress to pancreatic intraepithelial neoplasia (PanIN) and then pancreatic ductal adenocarcinoma (PDA). This is an example of Dvorak’s thesis, which states that cancer represents the ever-healing wound (1). The profound difference between the normal healing process and the perturbed response engendered by oncogene activation raises the important question of whether key cell types that result from ADM impact cancer progression. Here, we investigate the role of tuft cells as they appear in response to oncogene-induced metaplasia in a number of cancers of human significance including lung, stomach, intestine, and pancreas (2–6).

Tuft cells are solitary chemosensory cells found throughout the hollow organs of the respiratory and digestive tracts. They are readily identified by their striking morphological features, including long, blunt microvilli, deep actin rootlets, and an extensive tubulovesicular system in the supranuclear cytoplasm (7). Their expression of taste, neuronal, and inflammatory cell signaling factors is thought to enable monitoring of intraluminal homeostasis and to achieve local responses via effectors (8,9). Despite the discovery of this ‘peculiar cell’ by Jarvi and Keyriainen in 1956, little functional data existed until recently (10). Since 2016, several independent groups have demonstrated that intestinal tuft cells detect and combat parasite infection through expression of succinate receptor 1 (Sucnr1) and cytokine IL-25 (11–16). More recently, thymic tuft cells were shown to function in the development and polarization of thymic invariant natural killer T cells (17,18). These studies suggest that tuft cells play multifaceted roles in sensing and responding to diverse infectious and inflammatory conditions.

While tuft cells have been reported in rat and human pancreas, they are not normally present in the murine pancreas (5,19). Rather, we and others showed that tuft cells transdifferentiate from the acinar cell epithelium as part of ADM in response to oncogenic *Kras* expression (5,20). Interestingly, while they increase during the genesis of pancreatic intraepithelial neoplasia (PanIN), they are not detected in PDA (5,20). Tuft cell formation is also characteristic of human pancreatitis and PanIN, suggesting a conserved, but currently undefined role in early tumorigenesis (5).

Previous reports predicted tuft cell function in injury and oncogenesis based on marker expression. For example, tuft cells express high levels of tubulin kinase Dclk1, which is pro-tumorigenic in several organ systems and has been suggested to label long-lived progenitor cells in the pancreas and intestines (21,22). Dclk1 ablation in the intestine, however, does not eliminate tuft cell formation, implying that while it is one marker for tuft cells, it is not essential for their genesis (20,23). Dclk1 is also expressed in other cell types (see below), which would be collaterally affected by its deletion. Given the expression of Dclk1 in multiple cell types, and its lack of essentiality for tuft cell formation, we undertook a strategy to specifically eliminate tuft cells through genetic ablation of the tuft cell master regulator Pou2f3 (11). We show here that pancreas-specific ablation of Pou2f3 prevents pancreatic tuft cell formation. Surprisingly, and in contrast to expectations from prior work, tuft cell ablation accelerates tumorigenesis. This identifies a protective role for tuft cells during tumor progression. We investigated the mechanisms underlying tuft cell function using small cell number RNA-sequencing, ultrastructural analyses, and metabolic profiling. These studies produce a consistent picture involving tuft cell secretion of suppressive eicosanoids, such as prostaglandin D_2_ (PGD_2_), as an important mechanism of action for promoting pancreas homeostasis and restraining disease progression.

## Results

### Tuft cell ablation accelerates tumorigenesis

Transcription factor Pou2f3 (also known as Skn-1a or Oct11) is the master regulator of tuft cell formation in several organs and its expression has recently been identified in pancreatic tumorigenesis (11,24,25). We therefore used pancreas-specific Pou2f3 deletion as a genetic strategy to investigate tuft cell contribution to the formation of pancreatic intraepithelial neoplasia (PanIN) and tumor progression. The specificity of this strategy is supported by the fact that 99% of tuft cells in the *LSL-Kras^G12D^;Ptf1a*^*Cre*/+^ (*KC*) mouse model of pancreatic tumorigenesis, identified by co-expression of tuft cell markers Cox1 and acetylated α-tubulin, express Pou2f3 (790/800 cells, 8 mice) (Figure 1A). Conversely, 93% of Pou2f3+ cells are definitively tuft cells (744/800 cells, 8 mice).

**Figure 1.**
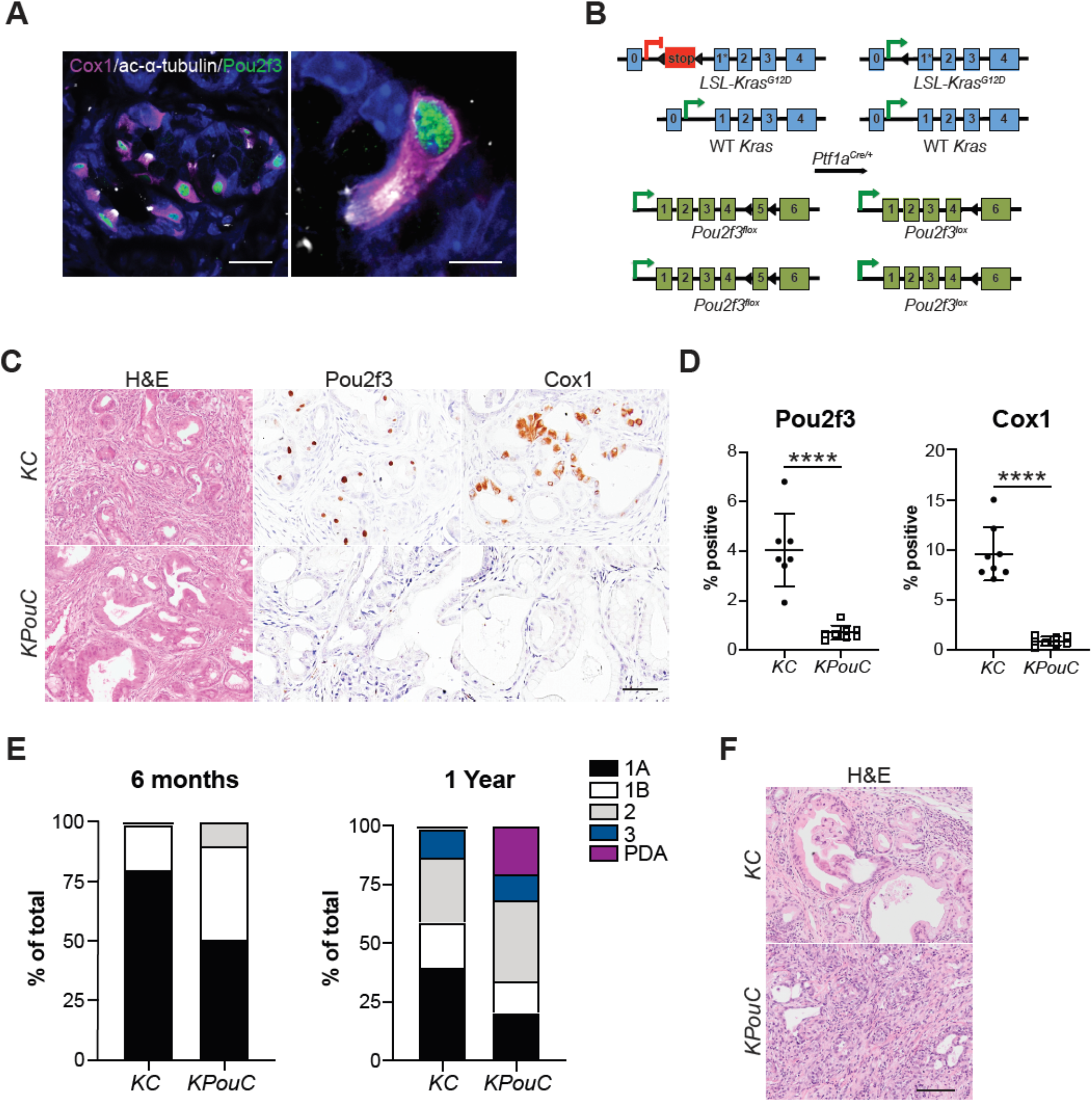
Pou2f3 and tuft cell ablation accelerates tumorigenesis. (**A**) Co-immunofluorescence for Pou2f3, green, and tuft cell markers Cox1, pink, and acetylated α-tubulin, white. Scale bars, 20 μm (left) and 5 μm (right). (**B**) Schematic of genetic modifications used to generate *KPouC* mice. Pancreas-specific Cre recombinase expression using *Ptf1a*^*Cre*/+^ results in removal of a stop cassette preceding a G12D mutation on exon 1 of the *Kras* allele as well as removal of exon 5 of *Pou2f3*. Arrowheads, loxP sites. (**C**) H&E and histology of *KC* and *KPouC* mice for Pou2f3 and tuft cell marker Cox1. Scale bar, 50μm. (**D**) Quantification of Pou2f3 and Cox1 staining in (C). (**E**) PanIN grams of pathologist-scored H&Es from age-matched *KC* and *KPouC* mice. (**F**) H&E of 1 year old *KC* and *KPouC* pancreata. Scale bar, 100μm. ****, p < 0.001.

Previous studies of tuft cell function used full-body Pou2f3 knockout mouse models, which could generate phenotypes due to tuft cell-specific and/or collateral effects (11). To increase specificity and eliminate influences from Pou2f3 ablation outside the pancreas, we generated a floxed *Pou2f3* mouse model enabling selective deletion in the pancreas (Figure 1B). Importantly, Pou2f3 is not expressed in the normal murine pancreas (Figure S1A). Consequently, pancreata from *Pou2f3^fl/fl^;Ptf1a*^*Cre*/+^ (*PouC*) and full body *Pou2f3*-/- mice were normal in size and contained intact exocrine and endocrine compartments with no overt pathology (Figure S1B-C). *Pou2f3^fl/fl^* mice were then bred into the *KC* model to generate *KPouC* mice. Age and sex matched *KC* and *KPouC* mice were sacrificed at 6 or 12 months and examined histologically. Pancreas-specific Pou2f3 deletion was confirmed by immunohistochemistry (IHC) (Figure 1C-D). We then confirmed tuft cell deletion using IHC for multiple markers (Cox1, Trpm5, and Vav1; Figure 1C-D, Figure S2A) as accepted tuft cell markers include inflammatory and neuronal proteins that are readily found in the stroma. Notably, Dclk1 was still expressed in *KPouC* pancreata, though at lower levels, demonstrating that its expression is not tuft cell-specific in the diseased pancreas (Figure S2A). Consistent with the pancreas specificity of this knockout model, we readily detected tuft cells and associated expression of Pou2f3, Dclk1, Cox1, Trpm5, and Vav1 in the intestines of *KPouC* mice (Figure S2B). Further, the lack of tuft cells in the pancreata of *KPouC* mice demonstrates that these cells do not migrate into the tissue during disease progression, but rather arise locally from the pancreas epithelium.

Although tuft cells are rare in the pancreata of 6-month-old *KC* mice, their absence in the *KPouC* pancreas correlated with accelerated tumorigenesis (Figure 1E-F). Pancreatic ductal adenocarcinoma (PDA) is thought to originate from metaplasia that has progressed through several steps of neoplasia (PanIN1a, PanIN1b, PanIN2, PanIN3) characterized by increasing nuclear atypia and loss of cellular polarity. Pathologist assessment of pancreata from 6-month-old mice by hematoxylin & eosin staining (H&E) from each cohort (n = 8 for each group) revealed significantly more PanIN1b (39% of lesions vs. 19%) and more PanIN2 (10% of lesions vs. 1.2%) in *KPouC* mice as compared to control (Figure 1E). By 12 months, 100% of *KPouC* mice (n = 3) had frank adenocarcinoma (PDA) whereas only a single cancerous lesion could be identified in one of the *KC* mice (n = 3) (Figure 1E-F).

Given the known expression of inflammatory regulators in tuft cells, we next investigated the impact of tuft cell loss on disease progression in the context of injury. To answer this question, 6-week-old *KC* and *KPouC* mice were given a short course of caerulein, to induce acinar cell injury, and were then allowed to recover for two weeks (Figure 2A). Pathologist scored H&E analysis revealed a trend towards higher grade PanIN in *KPouC* as compared to *KC* pancreata (Figure 2B-C). Immunohistochemical evaluation revealed significantly less normal tissue (amylase, 21% vs. 69%) and greater myofibroblast activation (αSMA, 41% vs. 27%) and extracellular matrix deposition (ECM, i.e. hyaluronan, collagen) in *KPouC* mice (Figure 2B, D, S3A-B). We identified significantly more apoptotic cells (CC3) and fewer proliferating cells (Ki67) as compared to *KC* pancreata (Figure S3A-B). There was a trend towards higher aberrant duct formation (Ck19) and immune cell infiltration (CD45) in *KPouC* mice, though it did not reach significance (Figure S3A-B).

**Figure 2.**
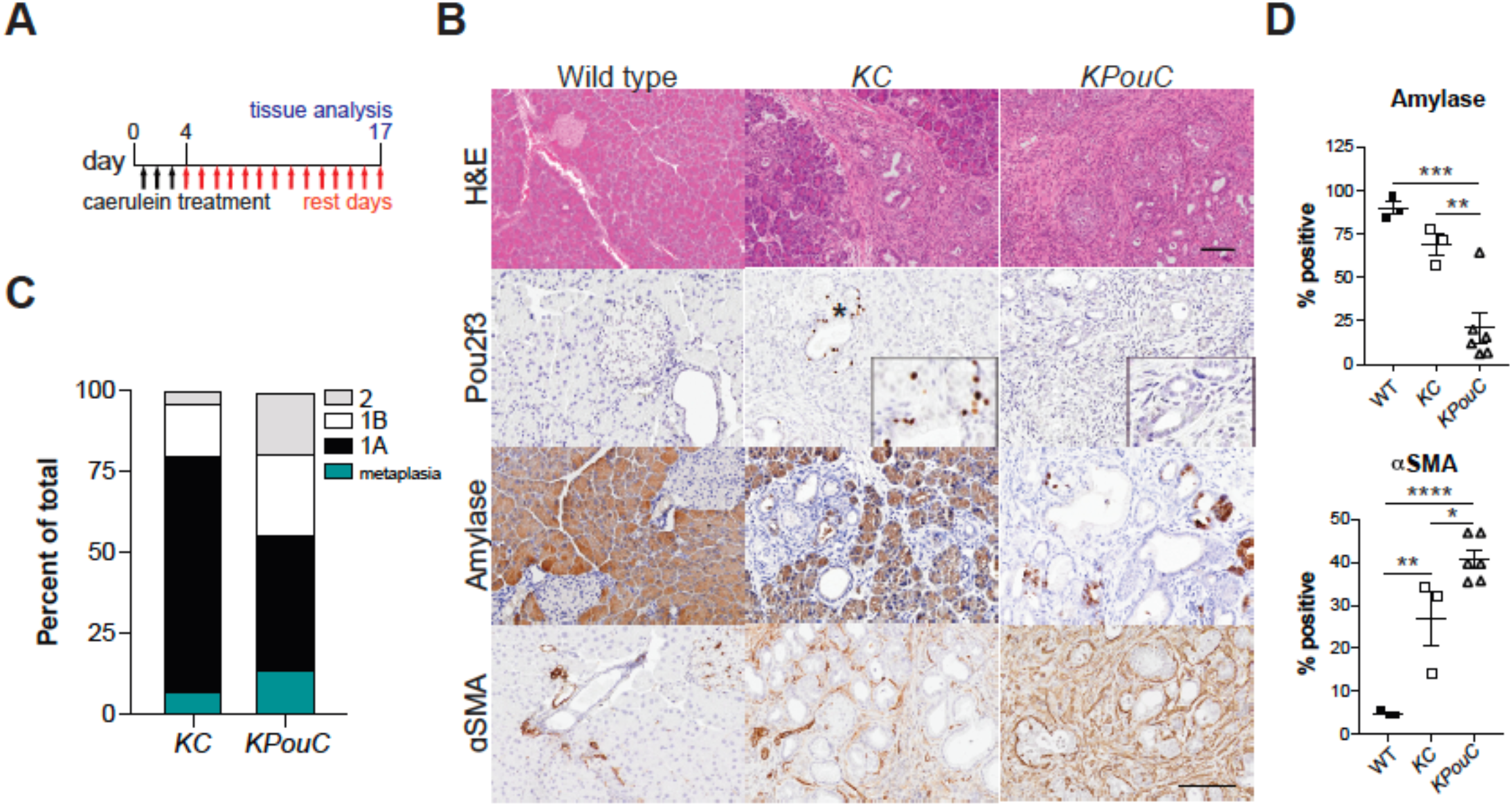
Tuft cell ablation enhances pancreatic injury. (**A**) Schematic for caerulein treatment of *KC* and *KPouC* mice. (**B**) H&E analysis and immunohistochemistry for Pou2f3, acinar marker amylase, and activated fibroblast marker αSMA, all brown. Scale bars, H&E and all histology panels, 100 μm; inserts, 50μm. (**C**) PanIN gram of pathologist-scored H&Es from treatment-matched *KC* and *KPouC* mice. (**D**) Quantification of histology shown in (B). *, p < 0.05; **, p < 0.01; ***, p < 0.005; ****, p < 0.001.

These observations indicate that while tuft cells are rare, they play an important role in pancreatic disease progression by moderating early responses to tissue injury. The data further identify an unexpected role for tuft cells in inhibiting pancreatic tumorigenesis. To determine the mechanism of tuft cell tumor suppression, we evaluated *KC* tuft cells transcriptomically and ultrastructurally to identify possible mechanisms of action.

### Transcriptomic analysis of tuft cells identifies lipid synthesis and metabolism pathways

We used RNA-sequencing to gain insight into potential mechanisms by which pancreatic tuft cells impact tumorigenesis. As the tuft cell population is rare, we used FACS for Siglec f to isolate enough cells to perform low cell number RNA-seq. While Siglec f labels Cd45+ eosinophils, it is also expressed in intestinal tuft cells (100%^+^, 158/158 cells, 3 mice) (9,11) and in pancreatic tuft cells in *KC* mice (99%^+^, 300/301 cells, 3 mice), but is not expressed in *KPouC* mice (Figure 3A, S2A). Using flow cytometry, we identified significantly more Siglec f+;EpCAM+ cells in 8-10 month old *KC* mice (~1.8% of the epithelium, range 1.0%-4.6%, n = 5) than in normal pancreas (0.001%, n = 5), consistent with histological studies (Figure. S4A) (5).

**Figure 3.**
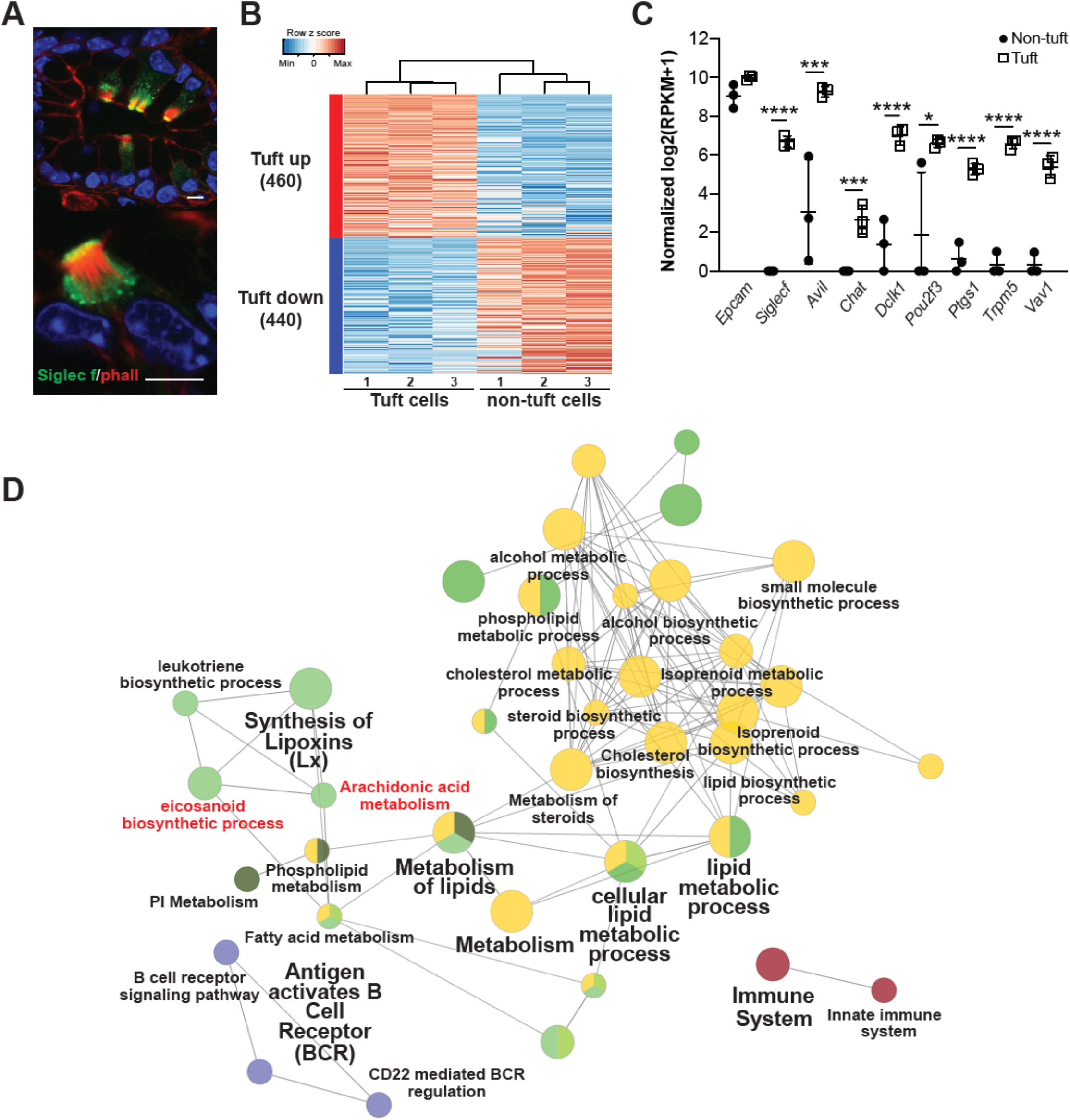
Transcriptomic analysis of *KC* tuft cells identifies lipid synthesis and metabolism pathways. (**A**) Co-immunofluorescence for phalloidin (red, labels the microvilli and actin rootlets of tuft cells) and Siglec f (green) in the *KC* pancreas. Scale bars, 5 μm. (**B**) Heat map with hierarchical clustering showing differentially expressed genes in Siglec f+;EpCAM+ tuft cells as compared to Siglec f-neg;EpCAM+ non-tuft epithelial cells. (**C**) Confirmation of Siglec f expression and tuft cell markers in sorted Siglec f+;EpCAM+ cells. *n* = 3 mice. (**D**) Gene ontology network analysis of genes differentially expressed in tuft cells identifies lipid synthesis and metabolism genes, notably eicosanoid biosynthesis-related genes, as enriched in tuft cells. *, p < 0.05; ***, p < 0.005; ****, p < 0.001.

We isolated 100 Siglec f+;EpCAM +;Cd45-neg tuft cells and an equivalent number of Siglec f-neg;EpCAM +;Cd45-neg non-tuft epithelial cells by FACS. We then isolated bulk RNA, prepared cDNA, and amplified it using the SmartSeq2 methodology to enable deep sequencing and comprehensive characterization of these small populations of cells (26). RNA-seq identified 900 genes differentially expressed between Siglec f+; EpCAM+;Cd45-neg and Siglec f-neg; EpCAM+;Cd45-neg populations (p<0.05 and average fold change>4) (Figure 3B). Tuft cell enrichment was confirmed in the Siglec f+ population by expression of markers such as *Pou2f3, Ptgs1* (Cox1), *Trpm5*, and *Vav1*, and low to no expression of acinar cell markers, such as *Ptf1a*, or islet cell markers, such as *Chga* (Figure 3C, S4B-C, File S1) (5,11,27). Furthermore, highly expressed genes in the Siglec f-neg population were enriched for gene sets associated with digestion, consistent with acinar and islet cell function (Figure S4D).

We used this small cell number sequencing approach to conduct RNA-seq on normal intestinal tuft cells to assess their similarity to pancreatic tuft cells. We found that both intestinal and pancreatic tuft cell signatures significantly overlap with single cell intestinal tuft cell sequencing data (Haber *et al*.,), validating our approach (Figure. S5) (28). To further validate the use of Siglec f to isolate tuft cells from PanIN, we sorted 116 Siglec f+; EpCAM +;Cd45-neg cells and conducted single cell sequencing by Smartseq2. We found significant enrichment of tuft cell markers identified by our bulk RNA-seq analysis in 88% (102/116) of these cells (FDR<0.05, File S1), confirming that Siglec f labels tuft cells in the pancreas (Figure S6).

Consistent with gene expression patterns identified in tuft cells from other systems, we identified expression of taste-signaling components (*Trpm5, Gnat3, Gng13, Itpr3*) and synaptic signaling markers (*Stx1a, Stx7, Snap23, Snap29, Vamp2, Nrgn, Gabra1, Gabbr1, Chat*) in *KC* tuft cells (File S1) (9,29,30). Importantly, and in contrast with data obtained from intestinal tuft cells, *Il25* and *Sucnr1* were not measurably expressed (Figure S7A,D, File S1). Due to the established functional role for these genes in intestinal tuft cells, we confirmed their absence by RT-qPCR (Figure S7B,E). Further, we crossed IL25-reporter mice (*IL25*^*F25*/+^) into the *KC* model and could not identify IL-25 expression in pancreatic tuft cells (Fig S7C). *Sucnr1* absence was confirmed with RNA *in situ* hybridization (Figure S7F). These data imply organ and/or context-specific tuft cell gene expression.

Network analysis of genes differentially expressed in *KC* tuft cells as compared to non-tuft epithelial cells highlighted a role in inflammation and identified pathways associated with lipid synthesis and metabolism, including eicosanoid biosynthesis and metabolism (Figure 3D). Eicosanoids are inflammatory lipid mediators with known roles in tumor progression. These data suggest that eicosanoid synthase Cox1 is not merely a tuft cell marker, but may also play a functional role.

### Lipid droplets serve as sites of eicosanoid synthesis in *KC* tuft cells

Tuft cells are known for their striking morphology. Therefore, to ascertain any structure-function relationships in pancreatic tuft cells, we conducted detailed ultrastructural analyses using scanning electron microscopy (SEM) and serial block face electron microscopy (SBFEM). SEM analysis of pancreata from year-old *KC* mice captured the topography of PanIN including the long, blunt microvilli characteristic of tuft cells (Figure 4A). Tuft cells (apical surface area = 27.84 + 7.55 μm^2^, 10 cells) were identified by microvilli (390 + 143 per cell, 10 cells) width (215.9 + 46.36 nm, 10 cells, 120 microvilli) and length (567.48 + 144.18 nm, 10 cells, 100 microvilli) (n = 2 mice). We also noted the possible budding of vesicles from tuft cell microvilli, suggesting an active secretory role for tuft cells in the PanIN stage of PDA progression (Figure 4A, S8A-B) (31).

**Figure 4.**
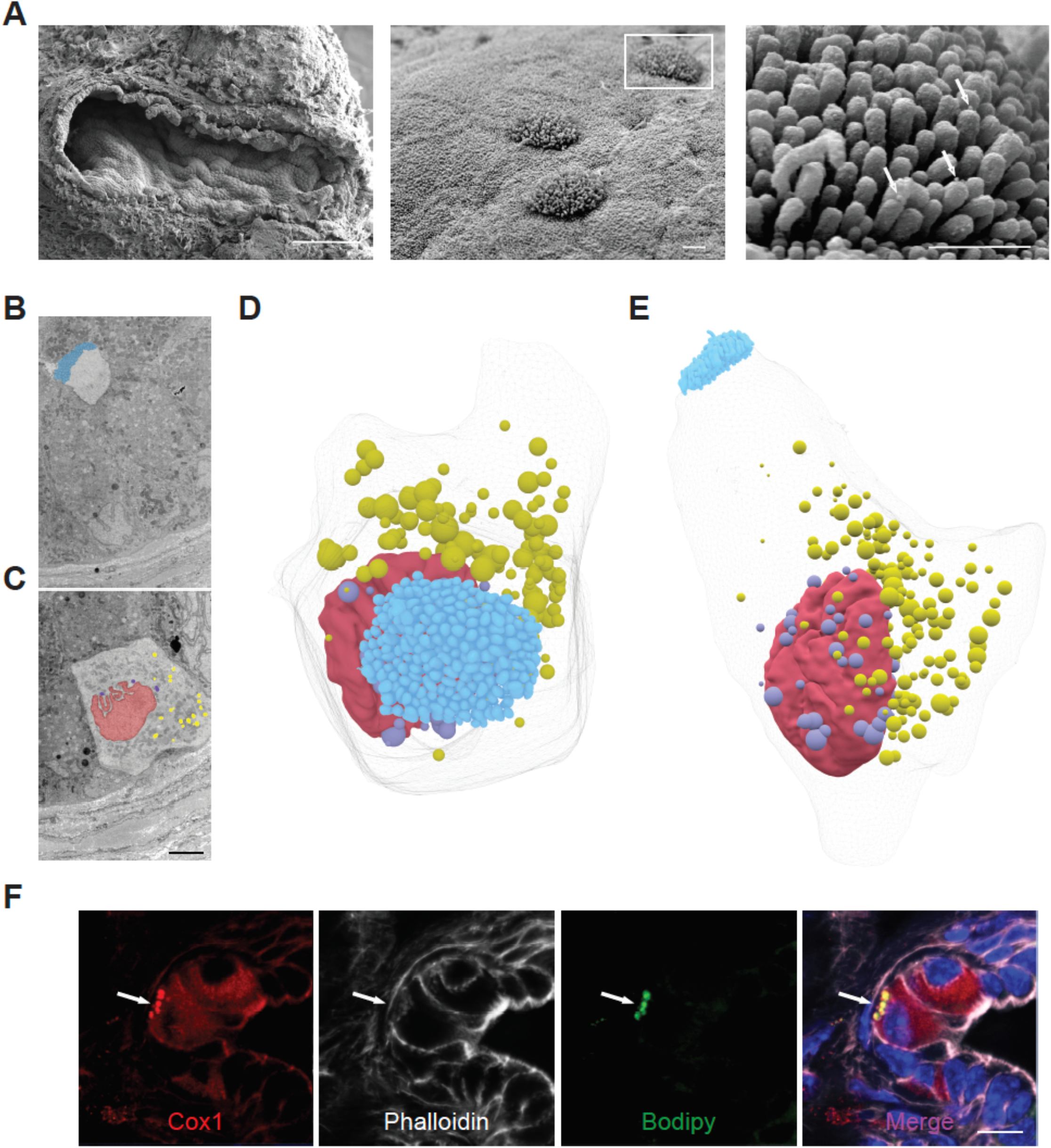
Lipid droplets as sites of eicosanoid synthesis in *KC* tuft cells. (**A**) SEM of PanIN from a year-old *KC* mouse (left panel, scale bar 100 μm) highlighting the tall, blunt microvilli of tuft cells (center panel), from which secretory vesicles appear to be budding (white arrows, right panel; center and right panel scale bars, 1μm). (**B**) Sections 45 and (**C**) 287 of 489, 100 nm sections through a single tuft cell captured by SBFEM. Scale bar, 5μm. (**D**) 3D reconstruction of all 489 of these sections highlighting the tuft cell microvilli (blue) and (**E**) nuclear-(lavender) and cytoplasm- (yellow) associated lipid droplets. Nucleus, pink. (**F**) Co-immunofluorescence for tuft cell marker and eicosanoid synthase Cox1 (12), phalloidin (50), and lipid droplet marker bodipy (green). Scale bar, 10 μm.

Identification of tuft cells by apical microvilli captured in a single plane obscures any information that may be gathered about peri-nuclear and basal structures. Recently, Hoover *et al*., used a combination of SBFEM and SEM imaging of serial ultra-thin sections to describe ultrastructural features of intestinal tuft in 3D (32). Therefore, to more completely evaluate pancreatic tuft cell ultrastructure, we performed SBFEM on PanIN from 6 and 12 month old *KC* mice. SBFEM uses an ultramicrotome mounted inside of a SEM to iteratively remove ultra-thin sections (<100 nm) of a resin embedded sample (or block) and images the newly revealed blockface with each slice. Images are produced by raster scanning the SEM beam across the sample surface and collecting the backscattered electrons that have interacted with the heavy-metal stained resin-embedded sample. We captured over 500 sections (100 nm thick) through a PanIN to reveal a nearly complete single tuft cell, capturing both the apical microvilli (Figure 4B) and the basal cytoplasmic constituents (Figure 4C) of the same cell. Segmentation and 3D reconstruction of some organelles from the processed image stack revealed the novel observation of over 180 perinuclear and cytoplasmic lipid droplets (Figure 4C-E, S8C-D, Movie S1). Lipid droplets are compact organelles lined by a phospholipid monolayer, which appear as dark smooth roughly spherical organelles when imaged by EM. Of 8 PanIN stacks acquired (~500 80-100 nm slices per PanIN), ~50% had tuft cells. In total, 9 tuft cells were analyzed and all had perinuclear and cytoplasmic lipid droplets (n = 3 mice), consistent with a functional role for lipid synthesis and metabolism in *KC* tuft cells (movie S2). We confirmed these EM observations by conducting co-immunofluorescence for tuft cell markers (Cox1, acetylated α-tubulin, and Dclk1) and the lipid marker Bodipy (Figure 4F, S9), resulting in detection of lipid droplets in 63% of *KC* tuft cells (188/300 cells, 3 mice). As lipid droplets may be obscured by the 2D nature of this analysis or may be below the visual detection limit, it is possible that they were also present in the other 37% of tuft cells.

Lipid droplets have traditionally been thought of as innocuous lipid storage organelles, but recent data emphasize their role as dynamic players in lipid metabolism and immune regulation (33). In fact, they have been reported to serve as inducible sites of eicosanoid synthesis in inflammatory cell populations, such as eosinophils, macrophages, and mast cells (34,35). In these populations, arachidonic acid is released from the lipid droplet phospholipid monolayer by cytosolic phospholipase A2 (cPLA2) and is then converted by Cox (cyclooxygenase) or Lox (lipoxygenase) enzymes to eicosanoids (33). Co-localization of Cox1 and Bodipy suggests that pancreatic tuft cell lipid droplets serve as sites of eicosanoid synthesis (Figure 4F). Interestingly, Cox1 localizes to the nuclear membrane in tuft cells and a number of lipid droplets were found to associate closely with the nucleus (Movie S2). Although mechanisms of tuft cell eicosanoid secretion have yet to be elucidated, the basal localization of lipid droplets and their contact with the dense tubulin network within tuft cells suggest that secretion may occur basally and/or apically (Figure S9).

### Eicosanoid and eicosanoid synthase expression in pancreatic neoplasia

Our RNA-seq analysis of *KC* tuft cells demonstrates significant enrichment of eicosanoid synthases, including lipoxin, leukotriene, and prostaglandin synthases as well as the entire pathway of proteins necessary to generate leukotriene LTC_4_ (Figure S10A-C). We determined which eicosanoids are present in pancreatic neoplasia by analyzing a comprehensive panel of 157 eicosanoids in whole tissue (pancreata) from either wild-type or 8-10 month old *KC*, PanIN-bearing mice by mass spectrometry. Prostaglandin D_2_ (PGD_2_) was present at extraordinarily high levels (369 +/- 102 pmol/mg, n = 5), while LTC_4_ and downstream products were absent (Figure 5A, File S2). We and others have documented expression of hematopoietic prostaglandin D_2_ synthase (Hpgds) in pancreatic tuft cells, which was confirmed here by RNA-seq (Figure S10B) (4,5,9). Correspondingly, we found Hpgds expression to be absent from the epithelium of *KPouC* mice by IHC, suggesting that epithelial Hpgds is tuft cell-specific (Figure S2A). Importantly, mass spectrometry on pancreata from 6 month old *KPouC* mice and age-matched *KC* mice from the same colony revealed a 72% decrease in PGD_2_ levels (695 +/- 296 vs. 195 +/- 164 pmol/mg) between the two genotypes (Figure 5B), suggesting that a significant amount of PGD_2_ derives from tuft cell-specific synthesis and secretion. It is noteworthy that the different genetically engineered mouse models (GEMMs) used in these experiments were raised in different backgrounds (methods) and were kept in different facilities and, thus, underwent tumorigenesis at different rates, requiring that controls for each experiment be taken from the same mouse colony.

**Figure 5.**
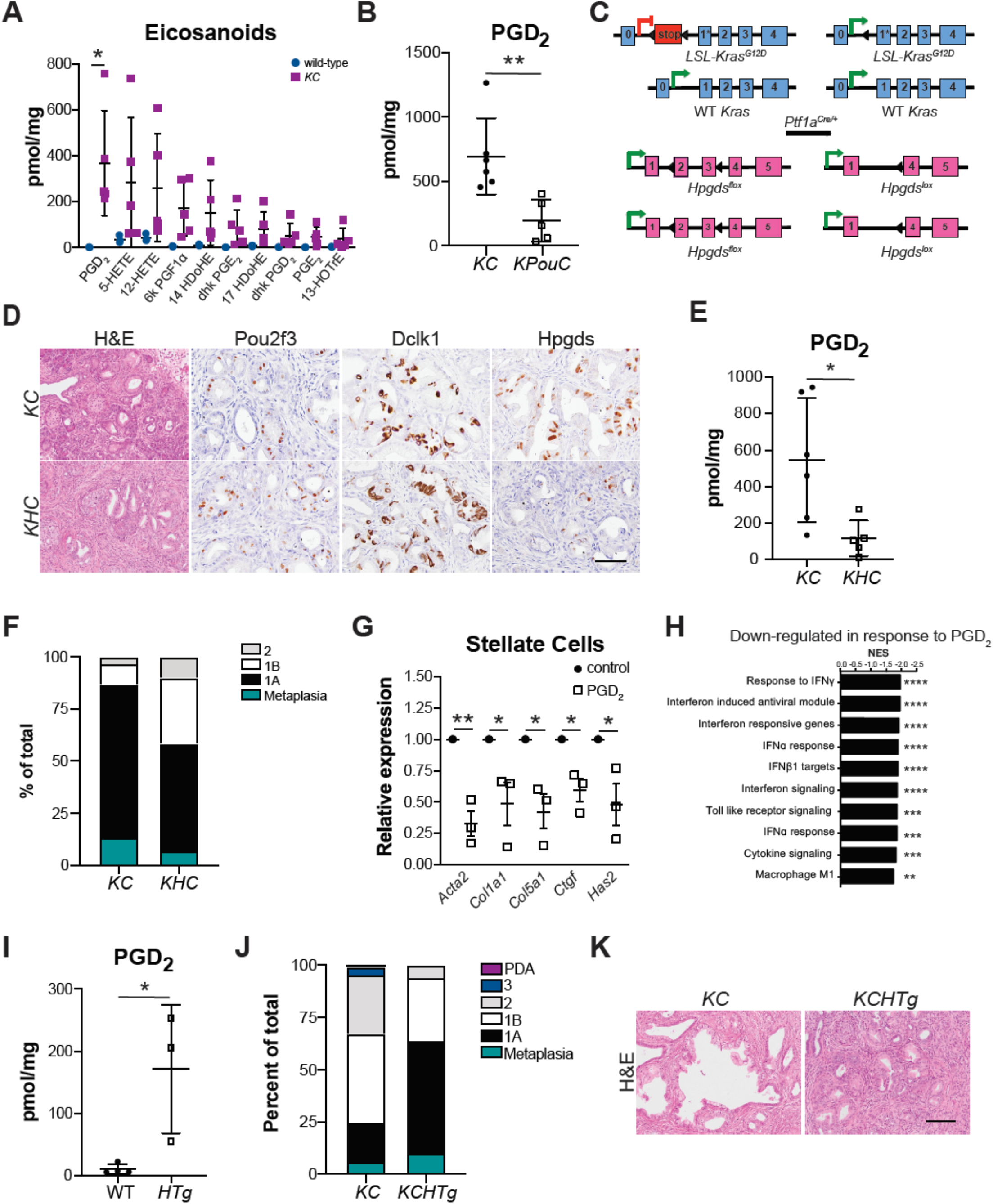
Tuft cell-derived PGD_2_ suppresses pancreatic tumorigenesis. (**A**) Eicosanoid expression profiling of whole pancreas tissue from 8-10 month old wild type or *KC* mice. (**B**) PGD_2_ levels in 6-month-old *KC* or *KPouC* mice. (**C**) Schematic of genetic modifications used to generate *KHC* mice. Pancreasspecific Cre recombinase expression using *Ptf1a*^*Cre*/+^ results in removal of a stop cassette preceding a G12D mutation on exon 1 of the *Kras* allele as well as removal of exons 2 and 3 of *Hpgds*. (**D**) Histological analysis of injury and disease progression in *KC* and *KHC* mice treated with caerulein. Pou2f3, Dclk1, and Hpgds, brown. Scale bar, 100 μm for H&E, 50 μm for immunohistochemistry. (**E**) PGD_2_ levels in caerulein treated *KC* or *KHC* mice. (**F**) PanIN gram of pathologist-scored H&Es from treatment-matched *KC* and *KHC* mice. (**G**) RT-qPCR of pancreatic stellate cells treated with control MeAOc or PGD_2_. (**H**) GSEA analysis of genes down regulated in LPS+PGD_2_ treated bone marrow macrophages as compared to LPS+MeAOc treatment. (**I**) PGD_2_ levels in 6-month-old wild type and *Hpgds-Tg* (*HTg*) mice. (**J-K**) PanIN gram and representative images of pathologist-scored H&Es from 6-month-old *KC* and *KCHTg* mice. Scale bar, 100μm. *, p < 0.05; **, p < 0.01; ***, p < 0.005; ****, p < 0.001; ns, not significant.

PGD_2_ has an extremely short half-life and has been shown to act locally (36). This begs the question of the relevance of epithelial, tuft cell-derived PGD_2_ to disease progression. We addressed this question by breeding a floxed *Hpgds^fl/fl^* allele into the *KC* model of pancreatic tumorigenesis to generate *KHC* mice (Figure 5C). This model selectively loses pancreatic epithelial Hpgds while maintaining stromal expression. Pancreata from *Hpgds^fl/fl^;Ptf1a*^*Cre*/+^ (*HC*) mice contain intact exocrine and endocrine compartments with no overt pathologies (Figure S11A). Six-week-old *KHC* (n=8) and control *KC* mice (n=6) from the same colony were given a short course of caerulein and were sacrificed two weeks later as previously described (Figure 2A). *KHC* mice treated with caerulein still form tuft cells, as identified by immunohistochemistry for markers Pou2f3, and Dclk1 (Figure 5D). Hpgds deletion was confirmed in *KHC* mice by IHC (Figure 5D), in situ hybridization (Figure S11B), and RT-qPCR of isolated epithelial cells collected by FACS (Figure S11C). To determine if deletion of epithelial-specific Hpgds is sufficient to decrease PGD_2_ levels, we conducted mass spectrometry on pancreata from caerulein treated *KHC* mice or treatment-matched *KC* mice from the same colony. We found a 79% decrease in PGD_2_ levels (543 +/- 341 vs. 114 +/- 99 pmol/mg) between the two genotypes (Figure 5E). Consistent with the phenotype determined in *KPouC* mice, pathologist-scored H&E analysis revealed significantly more PanIN1b (32% of lesions vs. 9.8%) and more PanIN2 (10% of lesions vs. 3.3%) in *KHC* mice as compared to control (Figure 5D, F). Collectively, these data demonstrate that tuft cell suppression of pancreatic tumorigenesis occurs, in part, through secretion of Hpgds-derived PGD_2_.

### PGD_2_ restrains disease progression by promoting environmental homeostasis

Elevated expression of stromal αSMA in *KPouC* mice, as compared to *KC* mice, suggests that loss of PGD_2_ may contribute to pancreatic stellate cell (PSC) activation and desmoplasia. We addressed this possibility by treating primary, activated murine PSCs with prostaglandins *in vitro*. Treatment with PGE_2_, as a proinflammatory control, increased expression of activation markers such as *Acta2* (αSMA) and growth factors such as *Ctgf* as determined by RNA-seq and confirmed by RT-qPCR (Figure S12A-C). By contrast, PGD_2_ treatment significantly decreased *Acta2* expression, as determined by RNA-seq and confirmed by RT-qPCR (Figure 5G, S12D-F, File S3). Correspondingly, we also saw a significant decrease in expression of ECM proteins *Col1a1* and *Col5a1, Ctgf*, and the hyaluronan synthase *Has2* (Figure 5G, S12E-F, File S3). Surprisingly, expression of PGD_2_ receptors *Ptgdr1* (Dp1) and *Ptgdr2* (Crth2/Dp2) was not detected by either RNA-seq or RT-qPCR in any sample (File S3, data not shown). However, PGD_2_ rapidly and non-enzymatically breaks down into a number of effectors, including variants of PGJ_2_, which are known ligands for nuclear receptor PPARγ. PPARγ was detected in mPSCs by RT-qPCR and, consistent with the literature, treatment of activated PSCs with the PPARγ agonist Rosiglitazone phenocopied PGD_2_ treatment (Figure S12G-H) (37).

Prostaglandins are known inflammatory mediators in injury formation, resolution, and tumorigenesis (38). PPARγ expression has been described in a number of immune cell populations and activation is known to suppress inflammation (39). As a model for assessing whether PGD_2_ itself is able to suppress inflammatory cell activation, we treated naïve or polarized bone marrow-derived macrophages (BMMs) *in vitro* with PGD_2_. RNA-seq analysis of naïve BMMs treated with PGD_2_ reveals induction of anti-oxidant genes and, as previously reported, a shift towards an anti-inflammatory macrophage phenotype (Figure S13A-B)(40). Lipopolysaccharide (LPS) induces a strong pro-inflammatory response in macrophages (Figure S13C), and subsequent treatment with PGD_2_ resulted in 322 genes being differentially expressed between control and LPS/PGD_2_-treated BMM (Figure S13D, File S4). PGD_2_ treatment resulted in suppression of proinflammatory interferon signaling pathways and a significant decrease in cytokine expression (Figure 5H, S13E-F).

We next turned to a genetic approach to determine whether PGD_2_ is sufficient to inhibit pancreatic tumorigenesis. We bred *HPGDS* transgenic mice, which express 375 fold more human *HPGDS* over endogenous mouse *Hpgds*, into the *KC* model (*KCHTg* mice) (41). Even with such a large increase in *HPGDS* expression, these mice are born with a normal pancreas with no overt pathologies (Figure S11D). Correspondingly, we found a significant increase in PGD_2_ levels in the pancreata of 6-month-old transgenic mice as compared to wild-type mice from the same colony (Figure 5I). Interestingly, when we examined PGD_2_ expression in 6-month-old *KC* and *KCHTg* mice, we did not identify significant differences in PGD_2_ levels, suggesting that mutant *Kras*-driven alterations to the pancreas are sufficient to drive high levels of PGD_2_ by 6 months of age (Figure S11E). However, when we examined disease progression, we found more advanced disease in 6-month-old *KC* mice (2/8 mice had frank adenocarcinoma) as compared to *KCTg* mice (0/7 mice had frank adenocarcinoma), suggesting that high PGD_2_ expression is sufficient to attenuate pancreatic tumorigenesis in early stages (Figure 5J-K).

### Pancreatic tuft cells in human disease

Tuft cells are also associated with human pancreatitis and PanIN, suggesting a conserved role across species (5). To infer the function of tuft cells in human pancreas disease, we evaluated tuft cell marker expression in RNA-seq data collected from human samples. To do this, we laser-capture dissected and sequenced the epithelium from 45 patients with precursor lesions (26 PanIN and 19 IPMN) and 197 patients with PDA (as described in (42). We then generated and overlaid a humanized version of our *KC* tuft cell signature on these data (KC Tuft single sample gene set enrichment analysis, ssGSEA) and found significantly higher expression in precursor lesions than cancer, consistent with tuft cell formation early in disease progression followed by loss later on (p<0.01, Figure 6A)(5). Tuft cell markers, such as *AVIL, TRPM5, GNG13* and *RGS13* were enriched in precursor lesions (Figure 6A) (9). Interestingly, and consistent with results in the *KC* mouse model, *IL25* was not detected. *HPGDS* expression, though, is significantly higher in precursor lesions than PDA, consistent with tuft cell expression (Figure 6B). To evaluate Hpgds protein expression, we conducted multiplex immunofluorescence for Hpgds and tuft cell marker phospho-EGFR on metaplasia and PanIN-bearing human pancreatitis samples (5). Though rare, we found that 100% of pancreatitis tuft cells (30/30; n=3 patients) express Hpgds at the protein level (Figure 6C), consistent with studies in normal human pancreas (19). We then conducted eicosanoid profiling on PDA (n=6) and nearby normal pancreas (n=3) (Figure S14A, File S5), but were unable to include human PanIN due to lack of available fresh tissue. Consistent with gene expression, PGD_2_ levels were very low in both PDA and nearby normal tissue (Figure S14B). Collectively, these data suggest that *Kras^G12D^*-induced tuft cells in both mouse and human disease utilize local, paracrine PGD_2_ signaling to curb activation of the tumor microenvironment and suppress disease progression (Figure 7).

**Figure 6.**
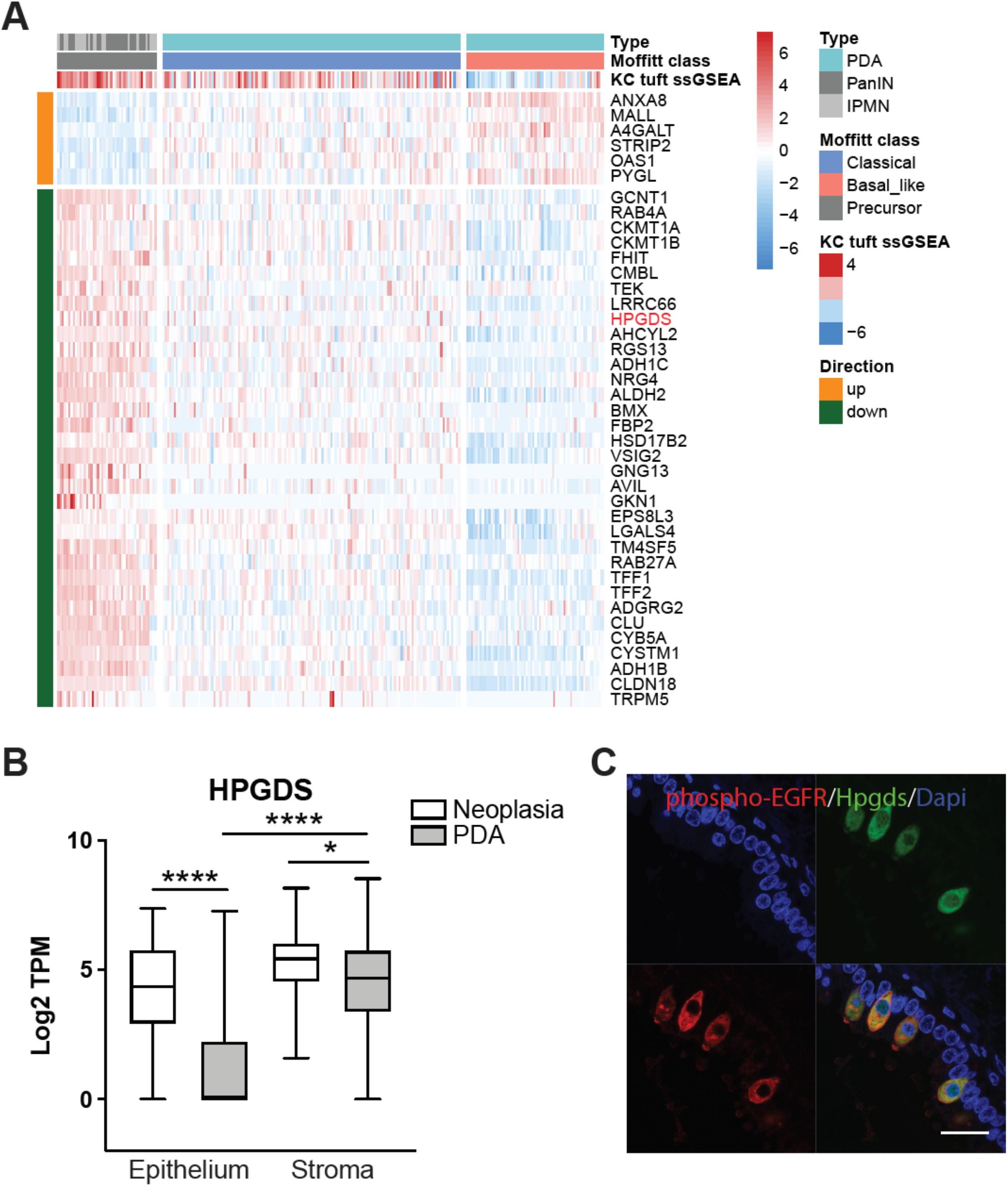
Tuft cell and Hpgds expression in human pancreatic tumorigenesis. (**A**) Heat map with hierarchical clustering showing a significant decrease in tuft cell-associated genes between pre-invasive PanIN and IPMN (n = 45) and PDA (n = 197), as well as between classical and basal-like PDA (p<0.01). ssGSEA, single sample gene set enrichment analysis. (**B**) RNA-seq of Hpgds in matched epithelial and stromal compartments from the same patients. (**C**) Co-immunofluorescence for tuft cell marker phospho-EGFR, red, and Hpgds, green, in human pancreatitis. Scale bar, 20μm.

**Figure 7.**
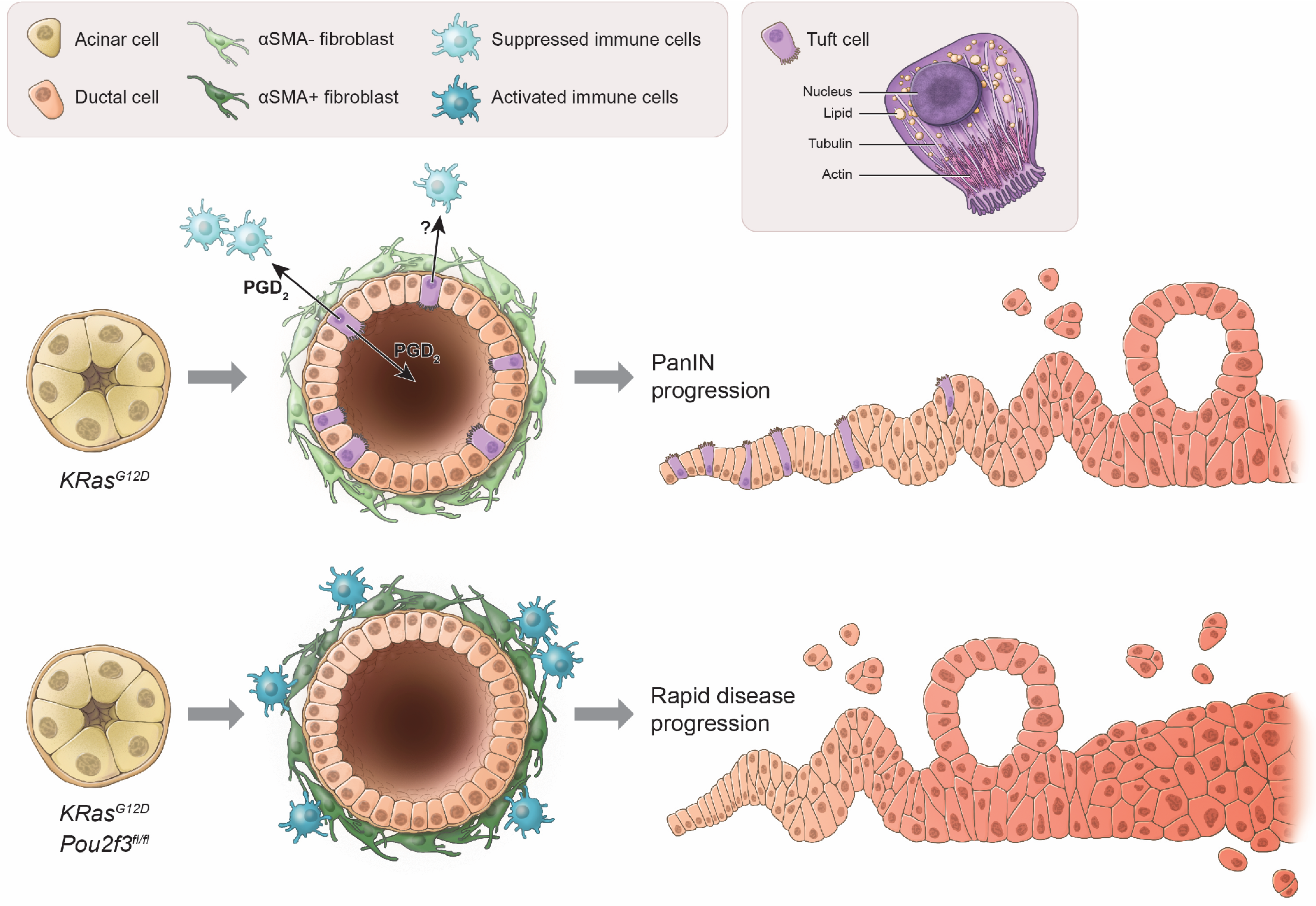
Model for tuft cell suppression of pancreatic tumorigenesis. The scheme at the top represents *Kras^G12D^*-induced pancreatic tumorigenesis. In this scenario, acinar cells expressing *Kras^G12D^* undergo metaplasia to form aberrant, heterogeneous, ductal structures containing tuft cells. Among the potential secretory products of tuft cells, PGD_2_ is released basally and/or apically and suppresses activation of associated stroma, including pancreatic stellate cells and immune cells. Metaplasia advances to pancreatic intraepithelial neoplasia (PanIN), but the process is slow. In the scheme at the bottom, acinar cells expressing *Kras^G12D^*, but depleted of Pou2f3, undergo metaplasia to form aberrant ductal structures lacking tuft cells. The lack of local, epithelial PGD_2_ secretion impairs homeostasis, leading to prolonged activation of stromal populations and accelerated pancreatic tumorigenesis.

## Discussion

Despite much conjecture over a pro-tumorigenic role for tuft cells in pancreatic tumorigenesis, we have found that eliminating tuft cells accelerates tumor formation, identifying a protective role for this cell type. Tuft cells, rather than seeding cancer, instill homeostasis in part through generation and secretion of lipid eicosanoid mediators. Consistent with this, our ultrastructural analyses identified the previously undescribed presence of lipid droplets in PanIN tuft cells, which may function in eicosanoid synthesis. Eicosanoid synthase expression, namely Cox1 and Hpgds, has been shown in tuft cells from a number of gastrointestinal organs and is often used to detect these cells (4,9). However, a functional role for tuft cell-derived prostaglandins has not previously been demonstrated. We show here that ablation of tuft cell PGD_2_ synthase Hpgds enhances pancreatic injury and disease progression. *HPGDS* and PGD_2_ overexpression is sufficient to inhibit pancreatic tumorigenesis. The global suppressive effects of PGD_2_ include inhibition of mPSC and BMM activation (Figure 7).

An anti-inflammatory role for PGD_2_ has previously been demonstrated in the context of injury and tumorigenesis in several organs (43). Correspondingly, reduction of PGD_2_ through Hpgds ablation has been shown to be pro-tumorigenic in lung and colon cancer models (41,43). Though we were unable to reliably detect either PGD_2_ receptor, we identified expression of derivative PGJ_2_ receptor PPARγ in both the epithelium and stroma. Within the epithelium, expression was largely restricted to tuft cells themselves suggesting an autocrine signaling loop. PPARγ has a known role in lipid droplet formation implying a role in eicosanoid synthesis with interesting implications for tuft cell regulation. Within the stroma, PPARγ activation is known to suppress mPSC activation and is a mechanism by which the inflammatory system regulates itself through suppression of Nf-κb (39,44). Further, PPARγ activation has been shown to inhibit tumor progression and metastases in mouse models of PDA (45).

While PGD_2_ synthesis and secretion is one attractive mechanism for tuft cell tumor suppression, these cells express additional immune modulators, including other eicosanoid synthases, which deserve further investigation (9). For example, in addition to Cox1, we have found that PanIN tuft cells express leukotriene synthase Alox5. While PGD_2_ is the most highly up-regulated eicosanoid in PanIN, Alox 5-derived 5-HETE is also highly expressed (Figure 5A) and has been described to have both pro and antiinflammatory roles. These data also underscore the importance of elucidating the role of eicosanoids in PDA formation and progression.

Interestingly, RNA-seq of PanIN tuft cells also identified a number of anti-bacterial and viral infection associated genes that could act in a pro-inflammatory manner (File S1)(46). These genes are also expressed in normal tuft cells from other organs, consistent with the predicted sentinel role for this cell type (9). While it is possible that these mediators are constitutively and simultaneously expressed, we favor a model where tuft cells function as a component of innate immunity and release different effectors depending on disease context. Consistent with this idea, we have discovered and characterized tuft cell formation in mouse models of pancreatitis (submitted manuscript). Though both are derived from acinar cells, pancreatitis and PanIN tuft cells differ significantly in expression of important effectors. For example, though we could not detect IL-25 in PanIN tuft cells, expression was evident in those associated with pancreatitis. These data demonstrate that we are just scratching the surface on understanding tuft cell function and regulation in disease.

These data also reinforce the view that metaplastic lesions function in tissue healing and injury resolution, but provide a new mechanistic perspective. Our data indicate that metaplastic lesions facilitate healing, in part, by generating differentiated, functional secretory cells, which signal to the microenvironment to contain disease progression, promote healing, and preserve organ function. In the setting of neoplasia, where the tissue cannot heal, we see an accumulation of tuft cells whose net effect is to abate transformation. To our knowledge, this is the first example of an epithelial-derived tumor suppressive cell type. Tuft cell formation may be one reason that humans can live with PanIN lesions for decades without progression to carcinoma. We often hear of mechanisms by which cancer cells evade the immune system and promote their own survival, but here we describe a mechanism by which the epithelium protects itself from these life-threatening events. Elucidating these protective mechanisms will allow us to understand when they fail or how to co-opt them for therapeutic benefit.

## Acknowledgments

We thank Eugene Ke, Alexis Roth, Annie Odelson, Hubert Tseng, and Gidsela Luna for technical expertise, Haiyong Han and Daniel Von Hoff for human samples, and Howard Crawford, Meggie Hoffman, Jakob von Moltke, Susan Kaech, and members of the Wahl laboratory for helpful discussions.

## Author contributions

Conceptualization: K.E.D. and G.M.W. Formal analysis: K.E.D., C.C., H.C.M., S.W.N., L.F., L.R.A., and C.O. Funding acquisition: K.E.D., G.M.W., U.M., and K.P.O. Investigation: K.E.D., R.R.G., S.W.N., R.F.N., W.H.A., C.T., C.R., and L.R.A. Project administration: K.E.D. Resources: H.C.M., M.R.S., M.O., Y.U., I.M., and K.P.O. Software: C.C., H.C.M., S.W.N., L.F., and C.O. Supervision: K.E.D., U.M., and G.M.W. Visualization: K.E.D., C.C., H.C.M., R.R.G., S.W.N., L.R.A., and C.O.

## Data and materials availability

Sequencing data that support the findings of this study are being deposited into GenBank and an accession code will be provided. Human RNA sequencing data may be found in GEO archive GSE110764. *Pou2f3^fl/fl^* mice are available from Dr. Matsumoto with a material transfer agreement.

## Materials and Methods

### Mice

Mice were housed in accordance with NIH guidelines in Association for Assessment and Accreditation of Laboratory Animal Care (AAALAC)-accredited facilities at the Salk Institute for Biological Studies. The Institutional Animal Care and Use Committee at the Salk Institute approved all animal studies (protocol 2011-0005). *LSL-Kras^G12D/+^; Ptf1a*^*Cre*/+^(*KC*, mixed C57BL/6J background) mice have been previously described in detail (47). *KC* mice conditionally express endogenous physiologic levels of activated *Kras^G12D^* targeted to progenitor cells of the developing pancreas. These animals spontaneously develop the full spectrum of precursor ductal lesions (pancreatic intraepithelial neoplasia, PanIN). FLARE25 (*Il25^F25/F25^*, C57BL/6J background) mice were generously provided by the Locksley (University of California San Francisco, CA) and von Moltke laboratories (University of Washington, WA)(13). *Pou2f3^fl/fl^* (*Pou*) mice were generated using a classical gene targeting method with BA1 embryonic stem cells, a hybrid of C57BL/6J and 129SvEv strains, by homologous recombination, resulting in deletion of 1303 base pairs from the *Pou2f3* gene. Chimeric mice were generated by inGenious Targeting Laboratories, were mated to flippase mice (in a C57BL/6J background), and then the obtained neo-deleted heterozygous mice were crossed with C57BL6J mice to remove the flippase allele. The resulting mice were pure heterozygotes (C57BL/6J, 87.5%; 129SvEv, 12.5%). *Hpgds^fl/fl^* (mixed C57Bl/6 strain background), and *Hpgds-Tg* mice (mixed FVB strain background) have been described previously (41,48).

### H&E scoring

For assessment of disease progression in 6-month-old *KC* vs. *KPouC* or *KC* vs. *KCHpgds-Tg* mice, 10, 10x images were scored from 2 x H&E slides, separated by a minimum of 200 μm for each mouse, by a licensed pathologist (V. Vavinskaya). For analysis of caerulein-treated *KC* vs. *KPouC* or *KC* vs. *KHC* mice, 10, 10x images were scored from 1 x H&E slide per mouse.

### Human pancreatitis samples

Distribution and use of all human samples was approved by the Institutional Review Boards of the Translational Genomics Research Institute and the Salk Institute for Biological Studies.

### Pancreatitis induction

Pancreatitis was induced with caerulein (Bachem) administered intraperitoneally (IP). Wild-type, *LSL-Kras^G12D^;Ptf1a*^*Cre*/+^ (*KC*), *LSL-Kras^G12D^;Pou2f3*^*fl/fl*^;*Ptf1a*^*Cre*/+^ (*KPouC*), *LSL-Kras^G12D^;Ptf1a*^*Cre*/+^;*Il25*^*F25*/+^, or *LSL-Kras^G12D^;Hpgds^fl/fl^;Ptf1a*^*Cre*/+^ (*KHC*) mice were given 250 μg/kg caerulein, once a day, for three consecutive days and were allowed to recover for two weeks.

### Histological staining and quantification

Tissues were fixed overnight in zinc-containing, neutral-buffered formalin (Fisher Scientific), embedded in paraffin, cut in 5 μm sections, mounted, and stained. Sections were deparaffinized in xylene, rehydrated in ethanol, and then washed in PBST and PBS. Endogenous peroxidase activity was blocked with a 1:50 solution of 30% H_2_O_2_:PBS followed by microwave antigen retrieval in 100 mM sodium citrate, pH 6.0. Sections were blocked with 1% bovine serum albumin (BSA) and 5% normal goat or rabbit serum in 10 mM Tris (pH 7.4), 100 mM MgCl_2_, and 0.5% Tween-20 for 1hr at room temperature, followed by an avidin/biotin blocking kit (Thermofisher) per the manufacturer’s instructions. Primary antibodies were diluted in blocking solution and incubated overnight. Information on primary antibodies is provided in Table S1. Slides were then washed, incubated in streptavidin-conjugated secondaries (for rabbit or mouse antibodies, Abcam, for rat or goat antibodies, Vector, for HABP, ABC HRP kit, Vector) and developed with DAB substrate (Vector). For trichrome staining, slides were stained using a kit (IHC world) according to the manufacturer’s instructions. Hematoxylin and eosin (H&E) staining was done to assess tissue morphology. All slides were scanned and imaged on an Olympus VS-120 Virtual Slide Scanning microscope. For quantification of histology, 10, 10x fields per scanned slide were scored in a blinded fashion using the ImageJ/FIJI plugin Immunohistochemistry (IHC) Image Analysis Toolbox (49). A statistical color detection model was trained based on multiple regions of interest (ROIs) manually selected from desired color pixel regions from sample images for each stain using the IHC Toolbox plugin. Each image was color deconvolved using its corresponding trained model within the plugin and a new RGB image containing only the isolated color was automatically generated. The hematoxylin counter stain was deconvolved in a similar manner. Using ImageJ/FIJI, the desired color-isolated image and the counter stain-isolated image were binarized and staining area of the two was measured by counting the number of pixels of foreground (50). The percentage of signal was determined by dividing the stain area by the sum of the stain area and the counter stain.

### Fluorescence microscopy

Tissues were fixed for 3-4 hours in 4% paraformaldehyde, washed 3x with PBS and floated overnight in 30% sucrose. Tissues were then incubated in a 1:1 mixture of 30% sucrose and Tissue-Tek optimal cutting temperature compound (OCT, VWR) for 30 min, embedded in OCT and frozen at −80°C. 7 μm tissue sections were cut, permeabilized with 0.1% Triton X-100 in 10 mM PBS, and blocked with 5% normal donkey serum and 1% BSA in 10 mM PBS for 1 hour at room temperature. Tissue sections were stained with primary antibodies in 10 mM PBS supplemented with 1% BSA and 0.1% Triton X-100 overnight (Table S1). Sections were then washed 3 x 15 min in PBS with 1% Triton X-100, incubated in Alexa Fluor secondary antibodies and/or phalloidin (Invitrogen), washed again for 3 x 5 min, rinsed with distilled water, and mounted with Prolong Gold containing Dapi (Invitrogen). Immunofluorescence on paraffin-embedded tissues followed the immunohistochemistry protocol until the blocking step. Instead, tissues were blocked in the donkey serum block described above and then followed the protocol for fluorescence microscopy described here. Tissues were imaged on either a Zeiss 710 confocal microscope or a Zeiss 880 Airyscan Super-Resolution microscope.

### Multiplex immunofluorescence

Co-expression of Hpgds and phospho-EGFR in human pancreatitis (antibodies, Table S1) was determined using a Perkin Elmer Opal 4-color Manual IHC Kit (NEL810001KT) per the manufacturer’s instructions.

### In situ hybridization

*Sucnr1* expression was validated in situ using the RNAScope Multiplex Fluorescent V2 kit (Advanced Cell Diagnostics, Catalog number 323110). The probe design and protocol was followed per the manufacturer’s instructions with antigen retrieval boiling for 15 minutes and Protease IV incubation at 40°c for 30 minutes. Bacterial-RNA probes were used as a negative control. Images were captured on a Carl Zeiss 880 Airyscan Super-Resolution microscope using 40X/1.2NA W objective with 1.8X digital zoom. Expression of exons 2 and 3 of murine Hpgds was determined by using the BaseScope Detection Reagent Kit v2 - RED (Advanced Cell Diagnostics, Catalog number 323900). The probe design and protocol was followed per the manufacturer’s instructions with antigen retrieval boiling for 15 minutes and Protease IV incubation at 40C for 30 minutes. Species-specific positive (probe targeting common housekeeping gene, catalog number 701081) and negative (probe targeting bacterial gene DapB, catalog number 701021) probes were used as both a technical quality control check and a RNA quality control check. All slides were scanned and imaged on an Olympus VS-120 Virtual Slide Scanning microscope.

### Scanning electron microscopy

Pancreata from 12-month-old *KC* mice were freshly dissected, cut into 2 mm fragments and immediately fixed with 4% formaldehyde (EMS), 2.5% glutaraldehyde (EMS), 0.1 M sodium cacodylate buffer (pH 7.2) (EMS), and 2 mM CaCL_2_, for 2 hours at room temperature. The samples were washed for 3 x 15 min in the same buffer, and impregnated with two consecutive baths of 1% Osmium tetroxide (EMS) and 1% tannic acid (Sigma-Aldrich) for 1hr each, with 3 x 10 min washes in between each bath. Samples were dehydrated in a graded series of ethanol until absolute, critical point dried (Leica CPD300) and placed on aluminum stubs, sputtered coated with platinum (4 nm) (Leica EM SCD500). Tissues were imaged on a FEG-SEM Zeiss Sigma VP, operated at 5kV and were photographed at 2048 x 1536 pixels of resolution.

### Serial block face scanning electron microscopy

Fresh pancreas samples were cut into 1 mm fragments and immersion fixed in a modified Karnovsky’s fixative (51) (2% paraformaldehyde (EMS), 2.5% glutaraldehyde (EMS), 2 mM CaCl2 in 0.1 M sodium cacodylate buffer, pH 7.2) at 37°C for 1 hour and kept in the same fixative overnight at 4°C. Samples were then washed in ice cold 0.1 M sodium cacodylate buffer 3×5 minutes before further post-fixation, staining, dehydration, and infiltration according to Deerinck *et al*. (https://ncmir.ucsd.edu/sbem-protocol), except that the final Durcupan (EMS) resin infiltration was doped with 1% Ketjen black carbon nanoparticles (gift from Nobuhiko Ohno, NIPS, Okazaki, Japan) to make the resin more conductive under the electron beam (52). Semi-thin sections (0.5-1.5 μm) were collected and examined with Toluidine blue staining using a light microscope until a region with a number of PanINs was identified. The block face was then trimmed to a roughly cubic face with a side-length of 300μm using glass and diamond trimming knives (Diatome) and an ultramicrotome (Leica) so that stained tissue was exposed on all sides, and a shallow notch was trimmed in a corner for orientation. The sample was then cut free from the block with a razor blade, and mounted flat on an aluminum 3View pin (Gatan) using toothpicks and conductive silver epoxy (EMS) before curing overnight in a 70°C oven. Pins were then sputtered (Leica) with about 15 nm of palladium, ultrathin sections were collected and the block face polished using a diamond knife, before finally loading pins into the 3View system (Gatan) mounted in a Zeiss Sigma VP scanning electron microscope. Over 500 sections, cut at a nominal thickness of 100 nm, were imaged with rasters of 20,000 x 20,000 pixels (1.2 kV, 1.5μs dwell time) with a pixel size of 16 nm pixels, creating a field of view measuring approximately 340μm x 340μm.

### Image Processing

Raw stacks were rigidly aligned using the Linear Stack Alignment with SIFT plugin (https://imagej.net/Linear_Stack_Alignment_with_SIFT) in Fiji (NIH) and the region of interest was cropped into a substack. An affine alignment was performed using SIFT again, and a gaussian blur of 0.7 pixels was applied to reduce apparent shot noise to ease segmentation. The series was then loaded into Reconstruct software (53) and the basement membrane, along with the tuft cell plasma membrane, nucleus, microvilli, and lipid droplets were traced through the series by hand. The segmented data were then imported into Cell Blender (54,55), an extension of Blender software (2019), from which 3D meshes were generated and optimally smoothed using GAMer (https://arxiv.org/pdf/1901.11008.pdf). The 3-D models of the segmented organelles were rendered and analyzed in Blender using Neuromorph (https://github.com/NeuroMorph-EPFL/NeuroMorph).

All images (IHC, IF, EM) were digitally enhanced to edit the color, brightness and contrast levels using Zen (Carl Zeiss), ImageJ (Fiji), and/or Photoshop (Adobe) software.

### Stellate cell isolation and culture

Mouse pancreatic stellate cells (mPSCs) were isolated from the pancreata of wild-type CD1 mice as previously described (56). Pancreata from 2-3 mice were pooled prior to isolation. Briefly, pancreatic tissue was minced and digested with 0.02% Pronase (Roche), 0.05% Collagenase P (Roche), and 0.1% Dnase (Roche) in Gey’s balanced salt solution (GBSS, Sigma) at 37°C for 20 min. Digested tissue was then filtered through a 100 μm cell strainer (Fisher). Washed cells were resuspended in 9.5 ml GBSS containing 0.3% BSA (Sigma) and 8 ml of 28.7% Histodenz solution (Sigma). The cell suspension was layered beneath GBSS containing 0.3% BSA, and centrifuged at 1400 x g for 25 min at 4°C. Cells of interest were harvested from the interface of the Histodenz solution and the aqueous solution. Isolated mPSCs were washed with GBSS and re-suspended in DMEM (Fisher Scientific) supplemented with 20% FBS (Peak Serum), 100 mM Sodium Pyruvate (Life Technologies), 2 mM L-Glutamine (Life Technologies), 1x non-essential amino acids (Life Technologies) and antibiotics (penicillin 100 U/ml and streptomycin 100 mg/ml, Life Technologies). mPSCs were allowed to activate in culture and were then treated for 48 hrs with either prostaglandins (10 μM), control (methyl acetate, MeOAc), or Ppar agonists (Rosiglitazone, 100nm; GW7647, 1μM; GW501516, 100nM) in serum-free media before collection in Trizol (Thermofisher) for RT-qPCR or RNA-sequencing. A minimum of three biological replicates was analyzed per experiment.

### Macrophage isolation, differentiation, and culture

Bone marrow-derived macrophages (BMM) were generated from bone marrow extracted from the femurs of male CD1 mice as previously described (57). Briefly, femurs were extracted and cleaned. Both ends of the bone were removed with scissors and flushed with a 22-gauge needle containing extraction media (DMEM containing FBS) into a petri dish. Cells were washed twice with extraction media and incubated in BMM high media containing DMEM, FBS, sodium pyruvate, supernatant from L-929 cells (L-sup) and penicillin/streptomycin, which was changed day 3 post-isolation. To induce M1 polarization, macrophages were treated with 100 ng/ml LPS for 4 hours prior to control or prostaglandin treatment. To induce M2 polarization, macrophages were treated with IL-4 and IL-13 (both 20 ng/ml) for 4 hours prior to addition of prostaglandins (10 μM) or control (MeOAc). Cells were collected 24 hrs later in Trizol for RNA-sequencing.

### Tuft cell preparation and cell sorting

Pancreatic tuft cells were isolated from 3, 8-10 month old *KC* PanIN-bearing. The pancreas was quickly dissected, minced in 5 ml of DMEM with FBS and allowed to incubate for 2 min. Supernatant and fat were removed and pancreatic tissue was then incubated in 10 ml DMEM supplemented with 1 mg/ml collagenase I (Sigma), 1 mg/ml soybean trypsin inhibitor (Gibco), 1-1.5 mg/ml hyaluronidase (depending on tumor burden, Sigma), and 250 μl of DNAse I, shaking gently at 37°C for a maximum of 30 min. Digestion was monitored and tissue was further digested mechanically by pipetting. Digested tissue was passed through a 100 μm filter, washed with FACS buffer (PBS, 1 mM EDTA, 0.5% BSA), and incubated with ACK lysing buffer (Gibco) to remove red blood cells before staining for FACS.

Intestinal tuft cells were isolated as previously described (11). Briefly, the proximal 5 cm of the murine small intestine was dissected, flushed, cut longitudinally, and rinsed to remove luminal contents. Segments were incubated in a 37°C shaker for 20 min in 5 ml PBS containing 2.5 mM EDTA, 0.75 mM dithiothreitol (DTT), and 10 μg/ml DNAse I. Tissues were shaken vigorously for 30s, large pieces of intestinal wall were removed, and cells were spun down at 4°C, 1200 rpm, for 5 min. Supernatant was removed and cells were re-suspended in 5 ml HBSS with 1.0 U/ml Dispase and 10μg/ml DNAse I, shaking for 10 min. Digested cells were passed through a 100 μm filter and incubated with ACK lysing buffer.

Single cell suspensions were incubated on ice with mouse Fc receptor block (BD Biosciences, 1:200) followed by antigen-specific antibodies in FACS buffer. DAPI (molecular probes, 1:1000) and Annexin V (Biolegend, Pacific Blue conjugate at 1:200) were used to exclude dead and dying cells. Cells were labeled with Cd45 (Alexa Fluor 488), EpCAM (Alexa Fluor 647), and Siglec F (PE) (Biolegend, 1:200). Fluorescence Minus One (FMO) staining controls were included for gating populations of interest. Cells were FACS purified at the Salk Institute’s Flow Cytometry core facility on a BD Biosciences Influx cell sorter (100-μm size nozzle, 1 x PBS sheath buffer with sheath pressure set to 20 PSI). Cells were sorted in 1-drop Single Cell sort mode for counting accuracy; these were deposited directly into lysis buffer composed of DNase/RNase-free water, Triton X-100, and Ribolock (Thermo Fischer) in a 96 well plate.

### RNA extraction and RT-qPCR

mPSC or BMM were lysed in 1mL of Trizol (Life Technologies) and total RNA isolation was performed using an RNeasy Micro Kit (Qiagen, 74004) according to the manufacturer’s instructions. cDNA synthesis was carried out using iScript RT Supermix (BIORAD, 1708840), and RT-qPCR was performed using Power SYBR Green PCR Master Mix (Applied biosystems) on the ABI 7900 detection system (Applied Biosystems). Relative expression values were determined using the standard curve method. RT-qPCR was performed on bulk tuft cells and non-tuft epithelial cells from *KC* mice (100 cells per group) after the amplification step of the SmartSeq2 protocol (26). Results for mPSCs and *KC* cells were normalized to the housekeeping gene *Rplp0*. Results for BMM were normalized to *Hprt*. For RT-qPCR of exons 2-3 of murine *Hpgds, KC* or *KHC* pancreata were digested (as described for tuft cell isolation) and 100-200k EpCAM+ cells were isolated by FACS. RNA was isolated using RNeasy Micro Plus and Mini kits (Qiagen, 74034) and converted to cDNA using iScript RT Supermix. Quantitative real-time PCR was performed using a QuantStudio 5 Real-Time PCR System (ThermoFisher) by mixing cDNAs, SYBR Green PCR Master Mix (ThermoFisher, 4309155) and gene specific primers. Primer sequences are available in Table S2. *Hpgds* RT-qPCR data was normalized by the delta-delta-Ct method to housekeeping gene *Gapdh*.

### RNA-seq library generation, High-throughput sequencing, and analysis

Low input bulk RNA sequencing (RNA-seq) on intestinal and pancreatic tuft cells was performed using the Smart-Seq2 protocol as previously described (26). In brief, a minimum of three biological replicates, each with 100 tuft cells from an individual mouse, was sorted directly into 2 μl of Smart-Seq2 lysis buffer. Full-length cDNA was generated and size distribution representing RNA integrity was checked with Agilent TapeStation 4200 to ensure RNA quality. cDNA were then amplified with 18-22 PCR cycles, tagmentated with TDE1, and amplified again with 10 PCR cycles using a Nextera XT kit (Illumina FC-131-1096). The sequencing library purification was performed with AMPure XP beads (Beckman Coulter A63881), and 50 bp single-end sequencing was performed with Illumina HiSeq 2500. After quality check with FastQC, the fastq reads were mapped to the mm9 mouse genome using Hisat2 (58), followed by transcript assembly and quantification with Stringtie and Ballgown (59). Between samples total gene expression distribution was checked with boxplot to ensure similar transcriptome quality. None expressed genes were defined as RPKM variance across samples < 1 and were removed, followed with log2 transformation and quantile normalization with the R package preprocessCore. Differential expression analysis between tuft and non-tuft epithelial cells was performed with empirical Bayes shrinkage and moderated t-test using the Limma package and p values were converted into false discovery rates (FDR) using the Benjamini-Hochberg procedure (60). Differential expression heat map plotting and hierarchical clustering (with Euclidean distance and complete linkage) were performed with heatmap.2 in R. Pathway analysis was performed with GSEA (http://software.broadinstitute.org/gsea/index.jsp) using a ranked differential expression score and the MSigDB geneset database (61). The differential expression score was calculated as: −log_10_(FDR) X log_2_(fold change), which factors in both statistical confidence and the effect size. Gene ontology network analyses of differentially expressed genes were performed with ClueGO plug-in of Cytoscape (62,63).

For sequencing of mPSCs and BMMs, RNA quality was assessed using the Agilent TapeStation 4200 and RNA-Seq libraries were prepared using the TruSeq stranded mRNA Sample Preparation Kit v2 according to Illumina protocols. Multiplexed libraries were validated using the Agilent TapeStation 4200, normalized and pooled for sequencing. High-throughput sequencing was performed on the HiSeq 2500 system (Illumina). Image analysis and base calling were done with Illumina CASAVA-1.8.2. For mPSCs, data analysis was performed as described above, except that an additional batch effect removal step was performed on the normalized RPKM value using the R package SVA (64).

### Smart-Seq2 single-cell analysis

Reads were aligned to the mm10 mouse genome using STAR version 2.5.3a (65). Mapping was carried out using default parameters, filtering non-canonical introns and allowing up to 10 mismatches per read and only keeping uniquely mapped reads. Raw transcript counts were obtained using HTSeq version 0.9.1 (66). The union of all detected genes was compiled into a single expression table (156 cells and 28,002 genes). Cells with less than 500 unique genes and genes expressed in fewer than 3 cells were filtered using the Seurat R package (67) leaving 116 cells and 10,876 genes. Genes were log-normalized, z-scaled, and centered prior to dimensionality reduction and visualization. Before performing dimensionality reduction, a list of 3,105 most variable genes was generated using the *FindVariableGenes* function from the Seurat R package which calculates the average expression and dispersion for each gene, places these genes into bins, and then calculates z-score for dispersion within each bin. We then performed principal component analysis (PCA) over the list of variable genes and the first ten principal components were used for t-stochastic neighbor embedding (t-SNE). Heatmaps were generated using heatmap.2 R package. The tuft cell signature genes (FDR<0.05, log2FC>1 from Supplemental File 1) were intersected with the genes expressed in the scRNA-seq Tuft cells and the resulting 99 up-regulated and 20 down-regulated genes were used to color the t-SNE plot. To calculate significance of the tuft cell signature in each cell, the top 100 expressed genes were tested for significant overrepresentation of 109 genes upregulated in Tuft cells (FDR<0.05) using the hypergeometric distribution and setting the background to 10,876 expressed genes in the scRNA-seq Tuft cells. The −log_10_(FDR) was used to color the t-SNE plots indicating level of significance (102/116 cells at an FDR threshold of 0.05).

### Comprehensive eicosanoid panel

Eicosanoid profiling was conducted on flash-frozen pancreatic tissue from wild-type, caerulein-treated, or 8-10 month old *KC* mice. Tissues were homogenized in 1 ml of PBS containing 10% ethanol and 300 μl were extracted using strata-x polymeric reverse phase columns (88-S100-UBJ Phenomenex). Samples were taken up in 50 μl of 63% H_2_0, 37% ACN, 0.02% Acetic Acid, and 10 μl was injected into UPLC (ACQUITY UPLC System, Waters) and analyzed on a Sciex 6500 Qtrap mass spectrometer at the University of California, San Diego, Lipidomics Core as previously described (68). Samples were normalized to total protein content. PGD_2_ levels in *KC* vs. *KPouC, KC* vs. *KHC, KC* vs. *KCHpgds-Tg*, and wild-type vs. *Hpgds-Tg* mice were evaluated by performing direct metabolomics at UNMC as previously described (69).

### Analysis of tuft cell gene signatures in human PDA

Compartment-specific gene expression profiles of human IPMN (n =19), PanIN (n = 26) and PDA (n = 197) were generated using laser capture microdissection with subsequent RNA sequencing as described previously (42,70). These samples have not been previously published (manuscript in preparation). Raw counts from RNA sequencing were then normalized to account for different library sizes, and the variance was stabilized by fitting the dispersion to a negative-binomial distribution as implemented in the DESeq2 R package (71). Differential expression analysis was carried out using the limma R package between basal-like and classical PDA, basal-like PDA and precursor lesions and classical PDA and precursor lesions, respectively, and only those tuft cell genes achieving an FDR <= 0.05 at least once were retained. T statistics per gene from the three comparisons were integrated using their median and only those genes with an absolute integrated t statistic > 2 were kept for display in a heatmap using the pheatmap R package (http://CRAN.R-project.org/package=pheatmap). Single sample GSEA (ssGSEA) of the KC tuft cell signature genes in human PDA was carried out using the VIPER framework (72) yielding normalized enrichment scores per sample.

### Overlaying tuft cell signature genes from Haber et al

Single cell gene expression from gut epithelium as described by Haber *et al*. (28) were retrieved from the Broad Institute’s Single Cell Portal and only the full length RNA-Seq data were used to identify genes differentially expressed between tuft cells and all other cell types. Differential expression analysis was carried out using raw TPM (transcripts per kilobase million) and the Wilcoxon ranks sum test als implemented in the *FindMarkers* function from the Seurat R package (73). The top 100 tuft cell markers identified in this way were tested for enrichment on the intestinal and pancreatic *KC* tuft signatures, respectively, using GSEA (61).

### Statistical analysis

Statistical analyses, data processing and heatmap plotting were performed in R (https://www.r-project.org/) and/or Prism (GraphPad). Statistical significance was calculated by either two-tailed unpaired t-tests assuming equal variance or one-way ANOVA. Data are expressed as mean ± standard deviation.

---

**Figure S1.**
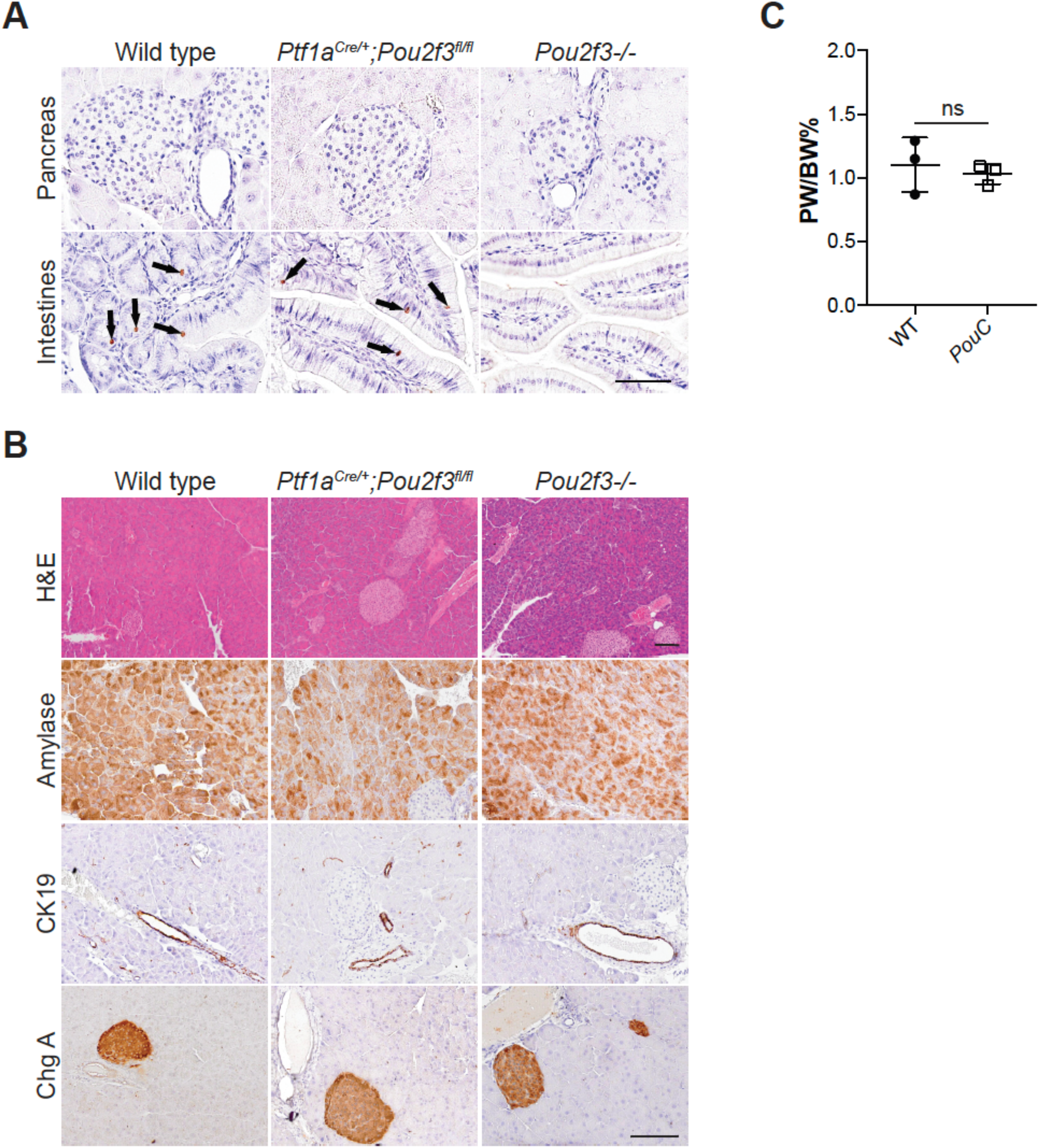
Pou2f3 ablation does not affect pancreas architecture or function. (**A**) Immunohistochemistry for Pou2f3 in the pancreata and intestines of wild-type, conditional *Pou2f3^fl/fl^;Ptf1a*^*Cre*/+^ (*PouC*), and full-body *Pou2f3*-/- knockout mice. Scale bar, 50 μm. (**B**) Histological analysis of pancreata from wild-type, *PouC*, and *Pou2f3*-/- mice. Amylase, acinar cells; Cytokeratin 19 (CK19), ductal cells; Chromagranin A (ChgA), islets. Scale bar, 100μm. (**C**) Pancreas:Body weight ratio (PW/BW%) of pancreata from wild-type and *PouC* mice. ns, not significant.

**Figure S2.**
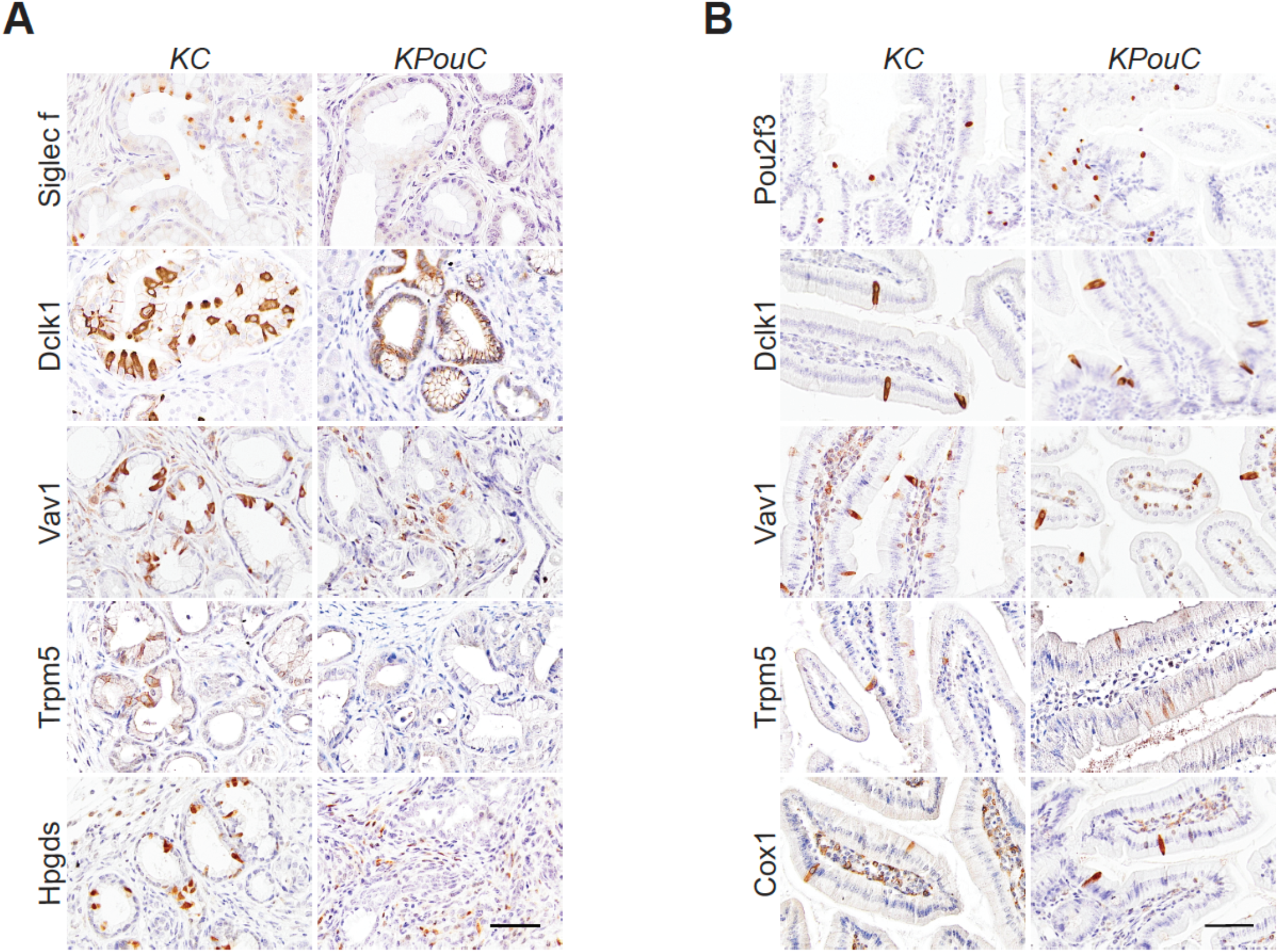
Pancreas-specific Pou2f3 ablation eliminates pancreatic, but not intestinal tuft cells. (**A**) Histological analysis of *KC* and *KPouC* pancreata for tuft cell markers Siglec f, Dclk1, Vav1, Trpm5, and Hpgds. (**B**) Histological analysis of *KC* and *KPouC* small intestines for Pou2f3 and tuft cell markers Dclk1, Vav1, Trpm5, and Cox1. Scale bars, 50 μm.

**Figure S3.**
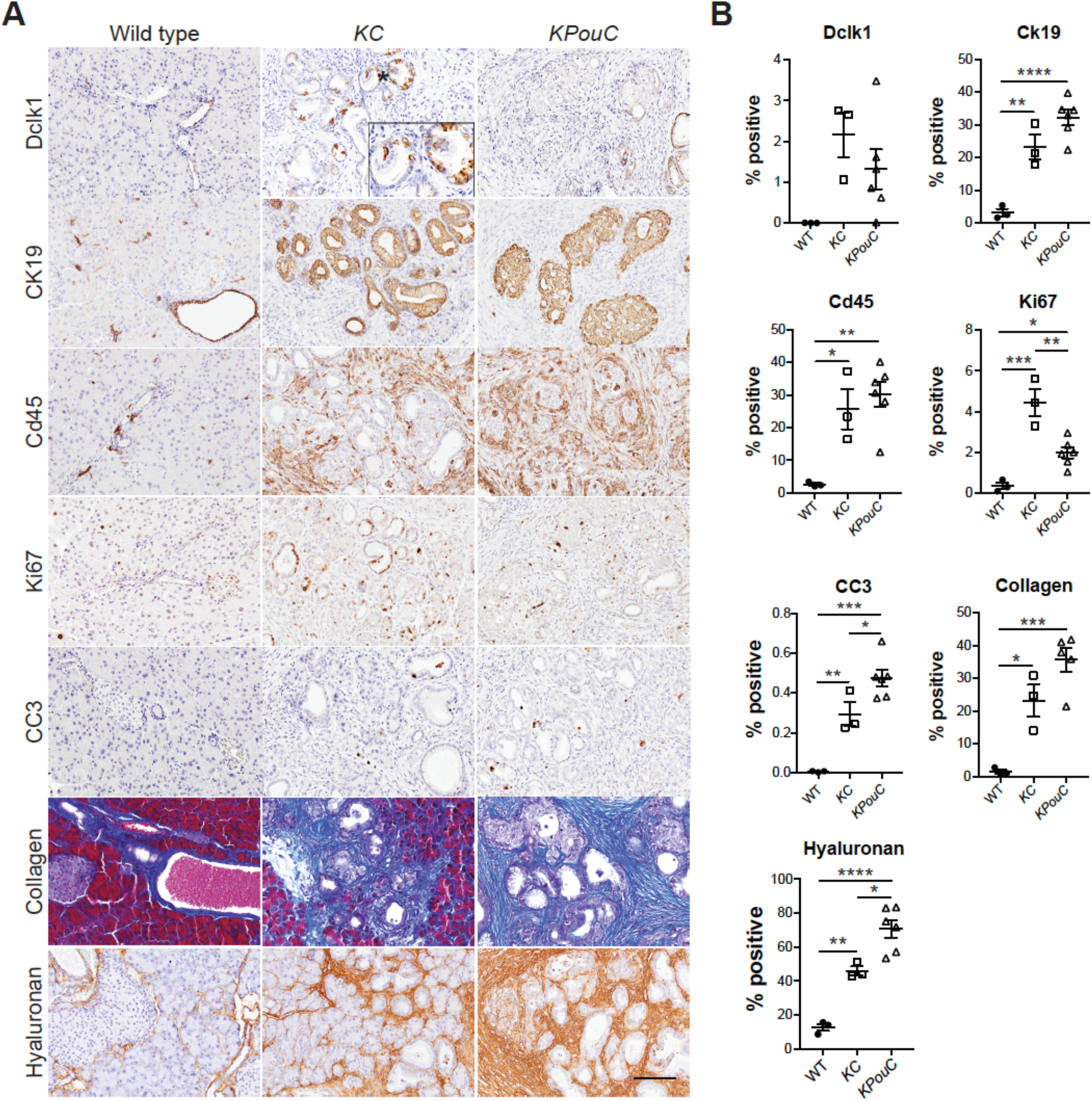
Histological analysis of disease progression in caerulein-treated *KC* and *KPouC* mice. (**A**) Immunohistochemistry for Dclk1, ductal cell marker CK19, inflammatory cell marker Cd45, proliferation marker Ki67, cell death marker CC3, extracellular matrix component hyaluronan (all brown), and collagen (trichrome, blue) in wild-type (WT), *KC*, and *KPouC* mice. Scale bar, 100μm. (**B**) Quantification of histology shown in (A). (WT, *n* = 3; *KC, n* = 3, *KPouC, n* = 6) *, p < 0.05; **, p < 0.01; *** p < 0.005; ****, p < 0.001.

**Figure S4.**
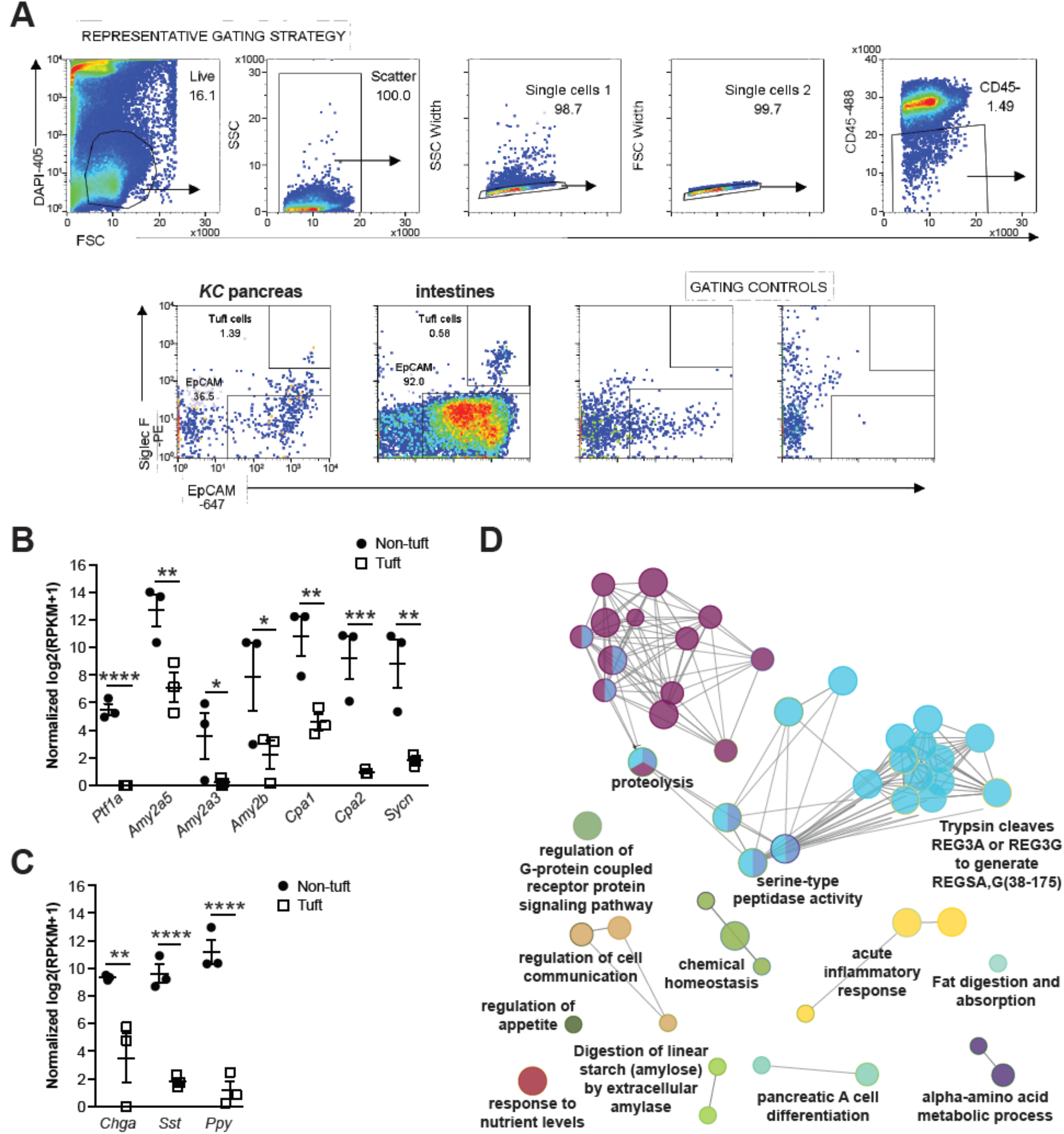
Fluorescence activated cell sorting of Siglec f+;EpCAM+ cells enriches pancreatic tuft cell populations. (**A**) FACS strategy for isolation of *KC* and intestinal Siglec f+;EpCAM+ tuft cells. (**B**) RNA-seq analysis demonstrates significantly lower expression of acinar and, (**C**) islet cell genes in Siglec f+;EpCAM+ sorted cells. *n* = 3 mice. (**D**) Gene ontology network analysis of genes significantly enriched in non-tuft epithelial cells identifies digestive pathways typically concentrated in pancreatic acinar and islet cells. *, p < 0.05; **, p < 0.01; *** p < 0.005; ****, p < 0.001.

**Figure S5.**
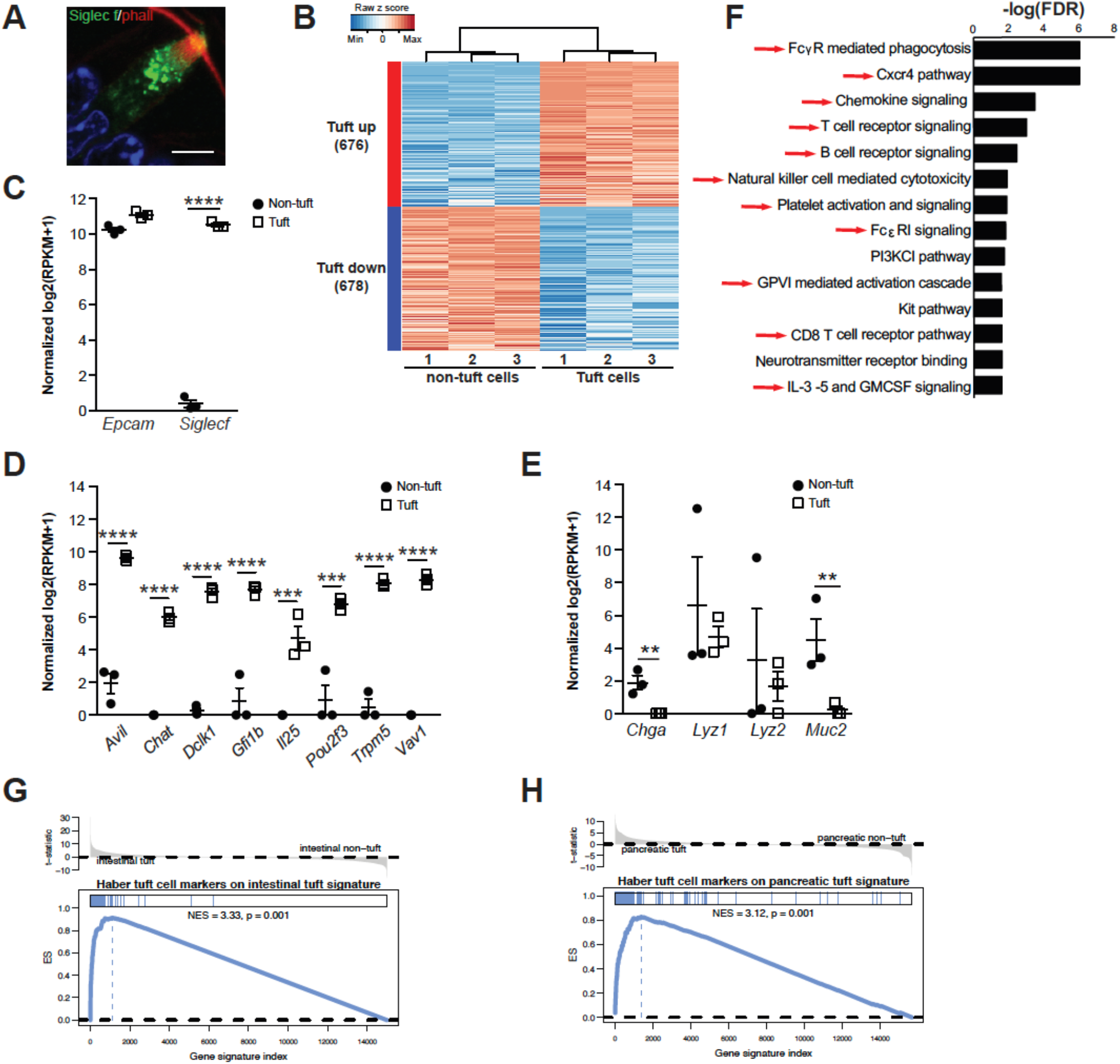
RNA sequencing and transcriptomic characterization of normal, intestinal tuft cells. (**A**) Co-immunofluorescence for Siglec f, green, and tuft cell marker phallodin, red. Scale bar, 5 μm. Image is representative of *n* = 3 mice. (**B**) Heat map with hierarchical clustering showing differentially expressed genes between Siglec f+;EpCAM+ tuft cells and Siglec f-neg;EpCAM+ non-tuft epithelial cells. (**C**) Confirmation of Siglec f expression and (**D**) tuft cell markers in sorted Siglec f+;EpCAM+ cells. (**E**) Exclusion of markers for other intestinal secretory cell populations. *n* = 3 mice for RNA-seq. (**F**) GSEA analysis demonstrating enrichment of inflammatory cell signaling pathway genes in intestinal tuft cells (red arrows). (**G**) Significant enrichment of tuft cell markers determined in Haber *et al*., in Siglec f+;EpCAM+ intestinal and (**H**) *KC* tuft cells. **, p < 0.01; *** p < 0.005; ****, p < 0.001.

**Figure S6.**
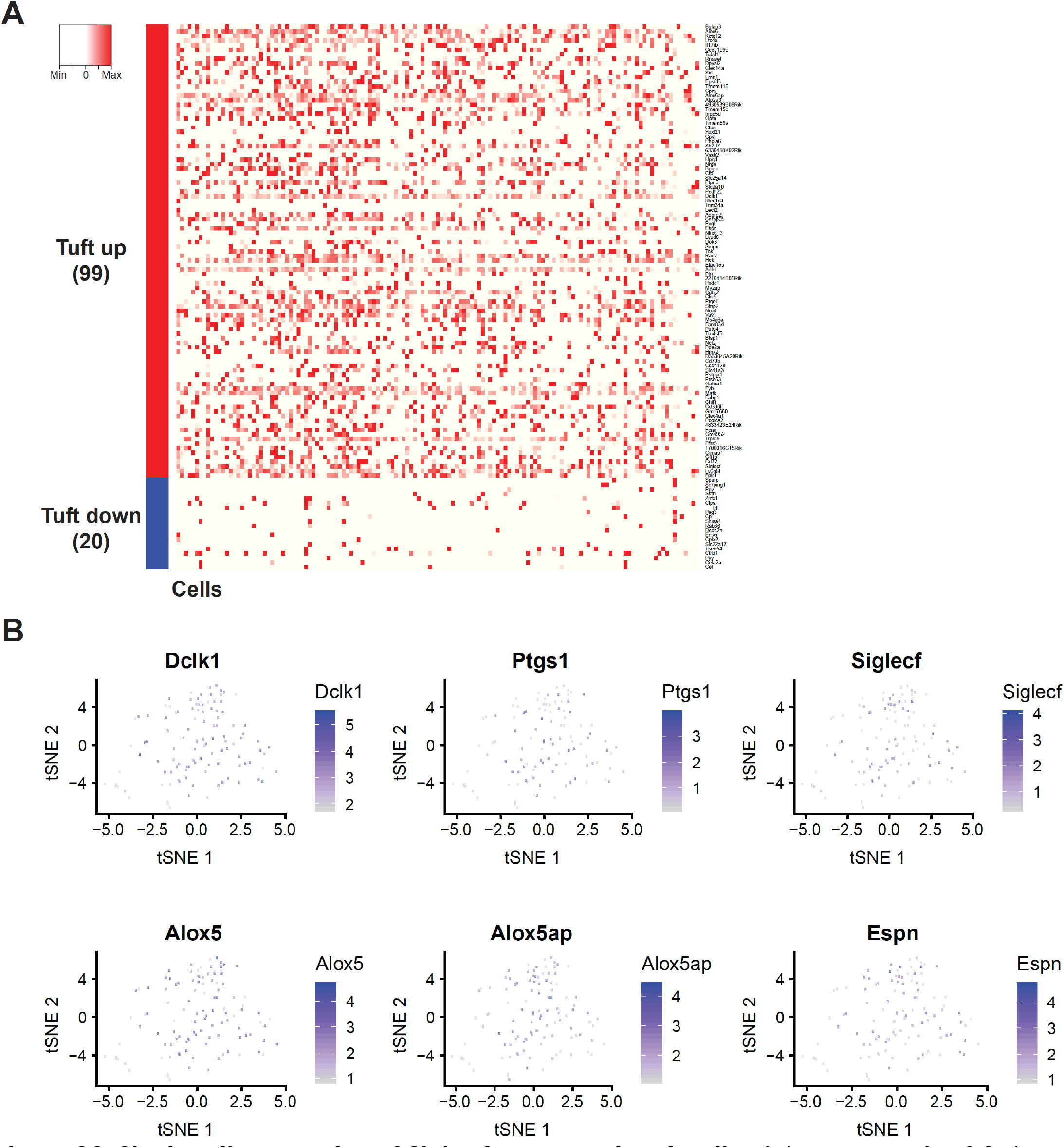
Single cell sequencing of Siglec f+ pancreatic tuft cells. (**A**) Heat map of 116 Siglec f+;EpCAM+ cells from 8-10 month old *KC* mice showing genes up and down regulated in tuft cells from bulk RNA-Seq. (**B**) t-stochastic neighbor embedding (t-SNE) of the same 116 cells showing high expression of selected tuft cell markers. Color intensity indicates the relative transcript level for the indicated gene in each cell.

**Figure S7.**
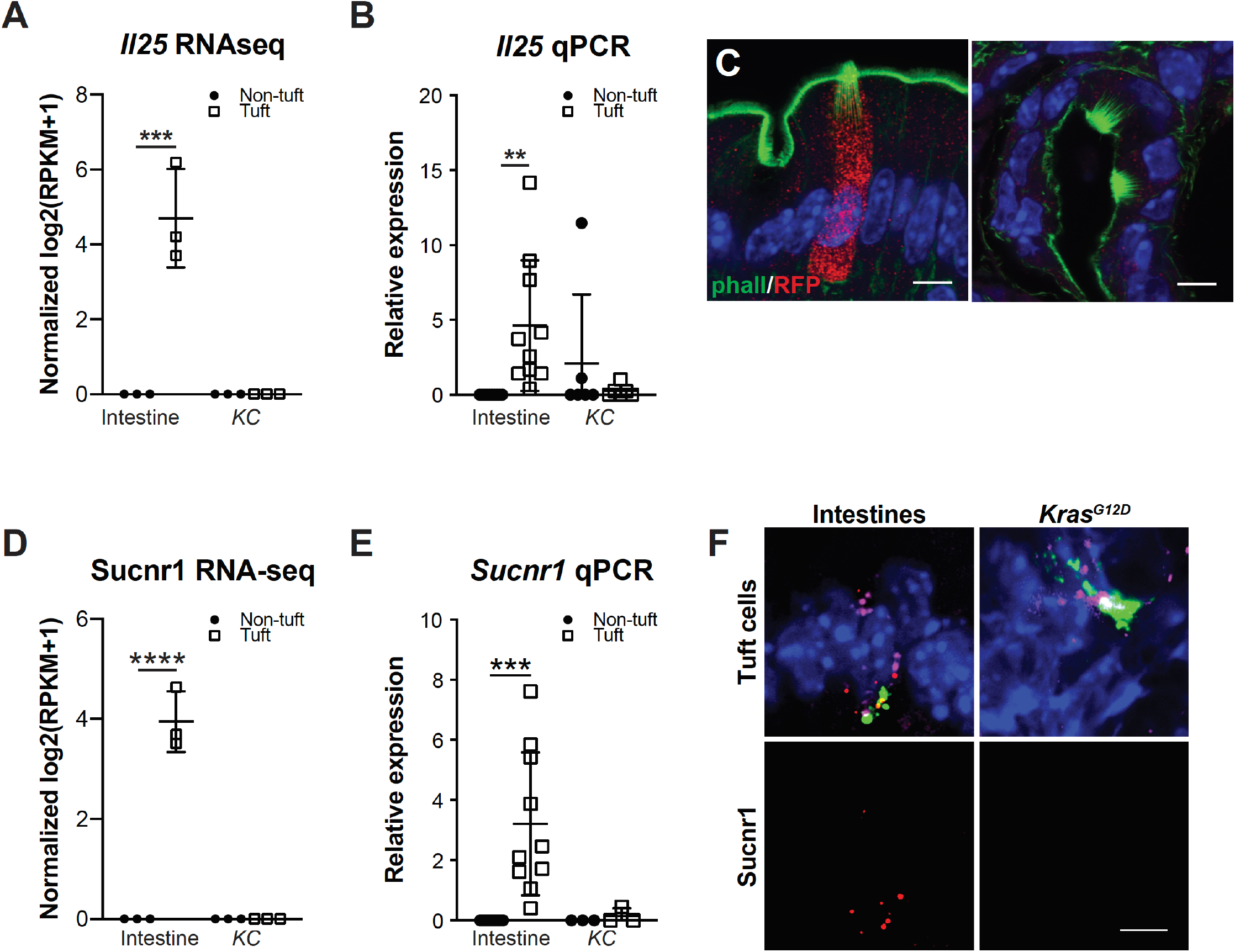
*Il25* and *Sucnr1* are not expressed in pancreatic tuft cells. (**A**) RNA-seq demonstrates *Il25* expression in intestinal, but not *KC* tuft cells (*n* = 3/group). (**B**) RT-qPCR confirms a lack of *Il25* in *KC* tuft cells (*n* = 6-9). (**C**) Co-immunofluorescence for red fluorescent protein (RFP, red) and phalloidin, green, in the intestines and pancreas of *Kras^G12D^;Ptf1a*^*Cre*/+^;*Il25^F25/F25^* mice. Scale bar, 5 μm. (**D**) RNA-seq for *Sucnr1* demonstrates expression in intestinal, but not *KC* tuft cells (n = 3/group). (**E**) RT-qPCR for *Sucnr1* confirms expression in intestinal, but not pancreatic tuft cells (n = 3-10). (**F**) RNA scope for *Sucnr1*, red, in tuft cells, identified by co-expression of Dclk1, green, and Cox-1 (*Ptgs1*, pink). Scale bar, 5μm. **, p < 0.01; ***, p < 0.005; ****, p < 0.001.

**Figure S8.**
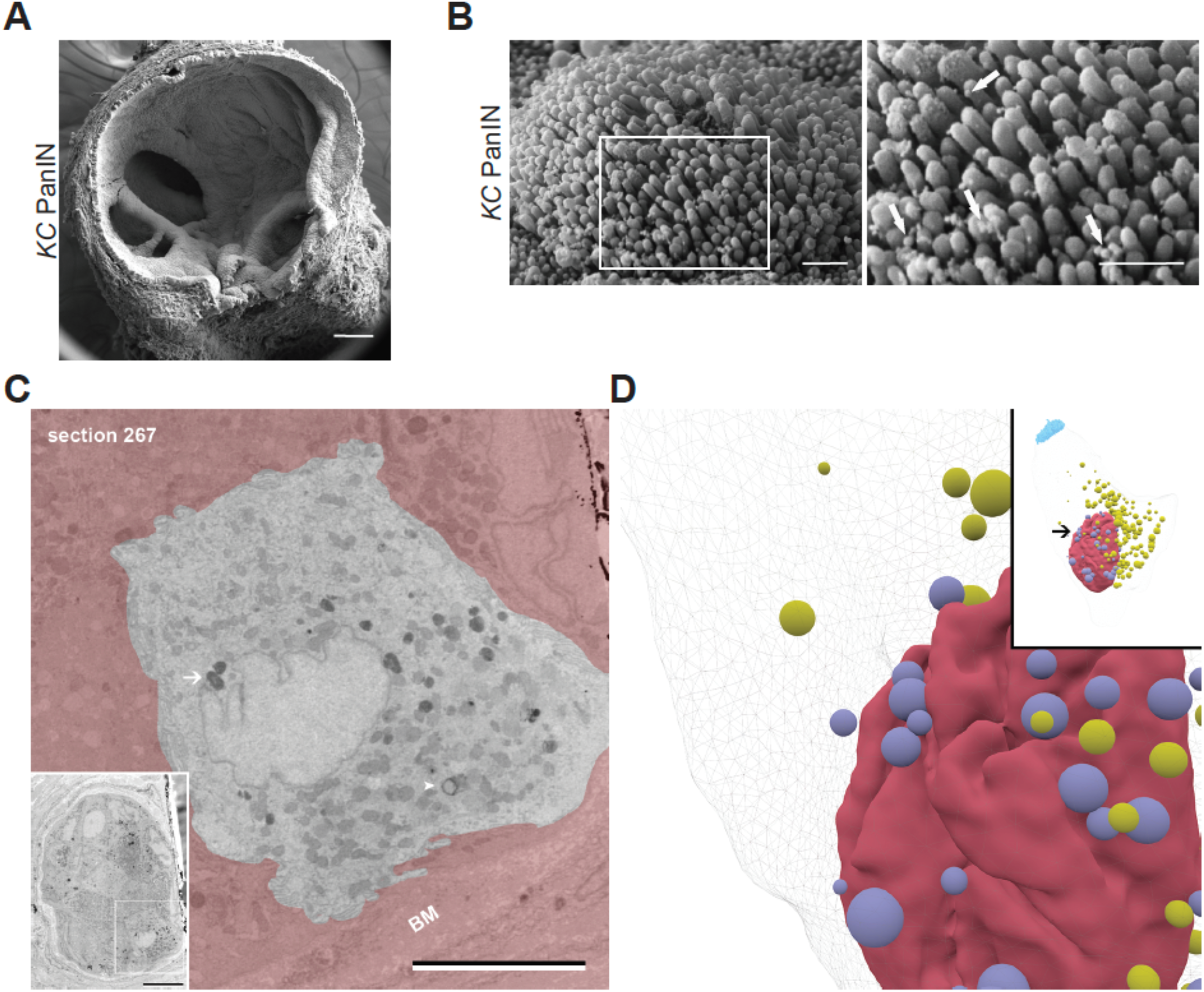
*Kras^G12D^*-induced pancreatic tuft cell morphology. (**A**) SEM analysis of a PanIN from a *KC* mouse. Scale bar, 100μm. (**B**) Vesicles (arrows) budding from the microvilli, imaged by SEM. Scale bars, 1μm. (**C**) SBFEM image (section 267) of a *KC* tuft cell highlighting nucleus- (white arrow) and cytoplasm-associated (white arrowhead) lipid droplets. BM, basement membrane. Scale bar, 5μm, inset 20μm. (**D**) Reconstruction of *KC* tuft cell lipid droplets highlighting those making contact with the nucleus, lavender. Cytoplasmic lipid droplets, yellow, nucleus, red, and microvilli, blue.

**Figure S9.**
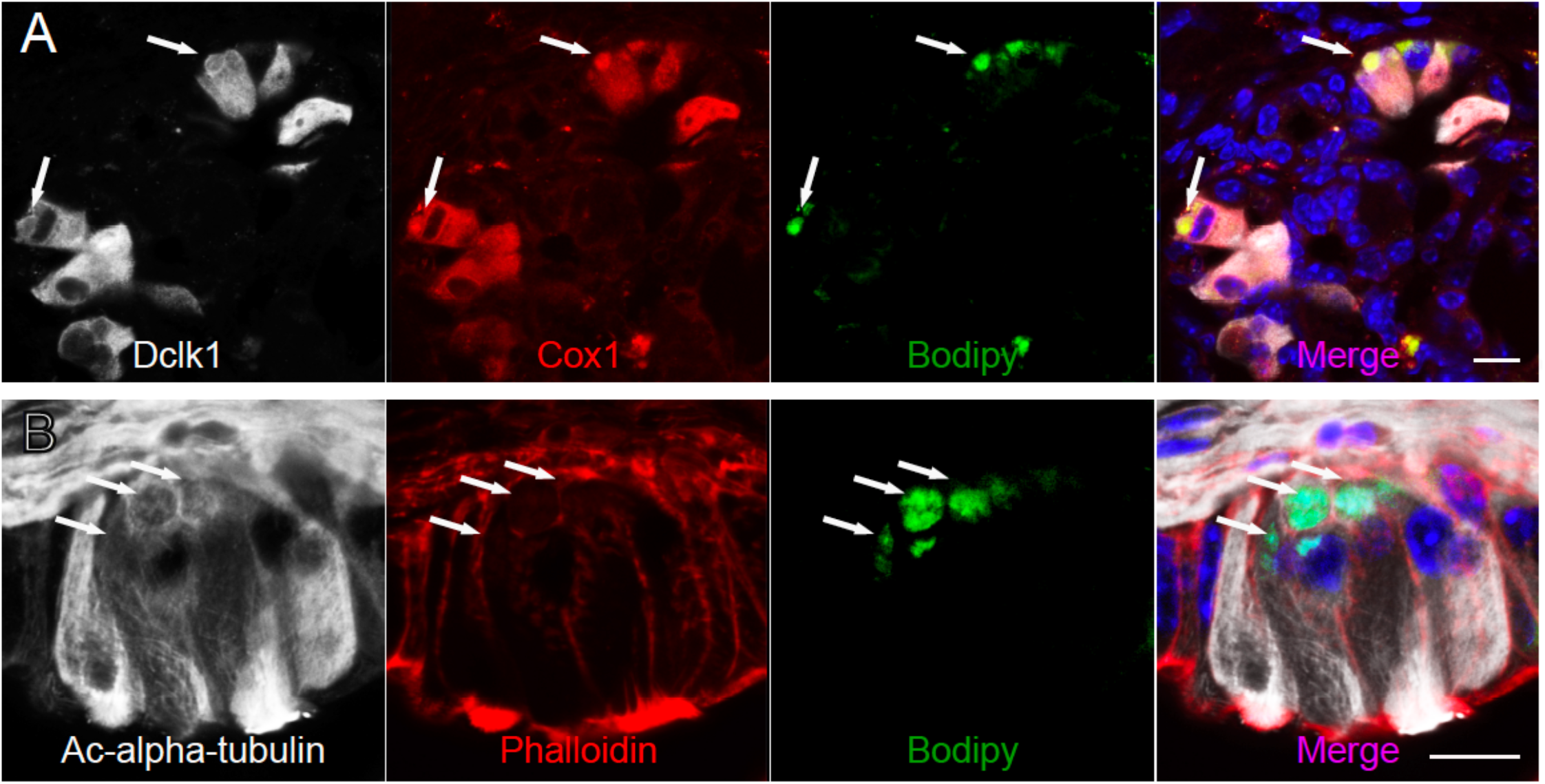
Lipid droplets as sites of *de novo* eicosanoid synthesis in *KC* tuft cells. (**A**) Coimmunofluorescence for tuft cell markers Dclk1, white, and Cox1, red, and lipid marker bodipy, green. (**B**) Co-immunofluorescence for phalloidin (red, marks the tuft cell microvilli and actin rootlets), acetylated alpha tubulin (white, marks the tuft cell dense tubulin network), and bodipy (green). Scale bars, 10 μm.

**Figure S10.**
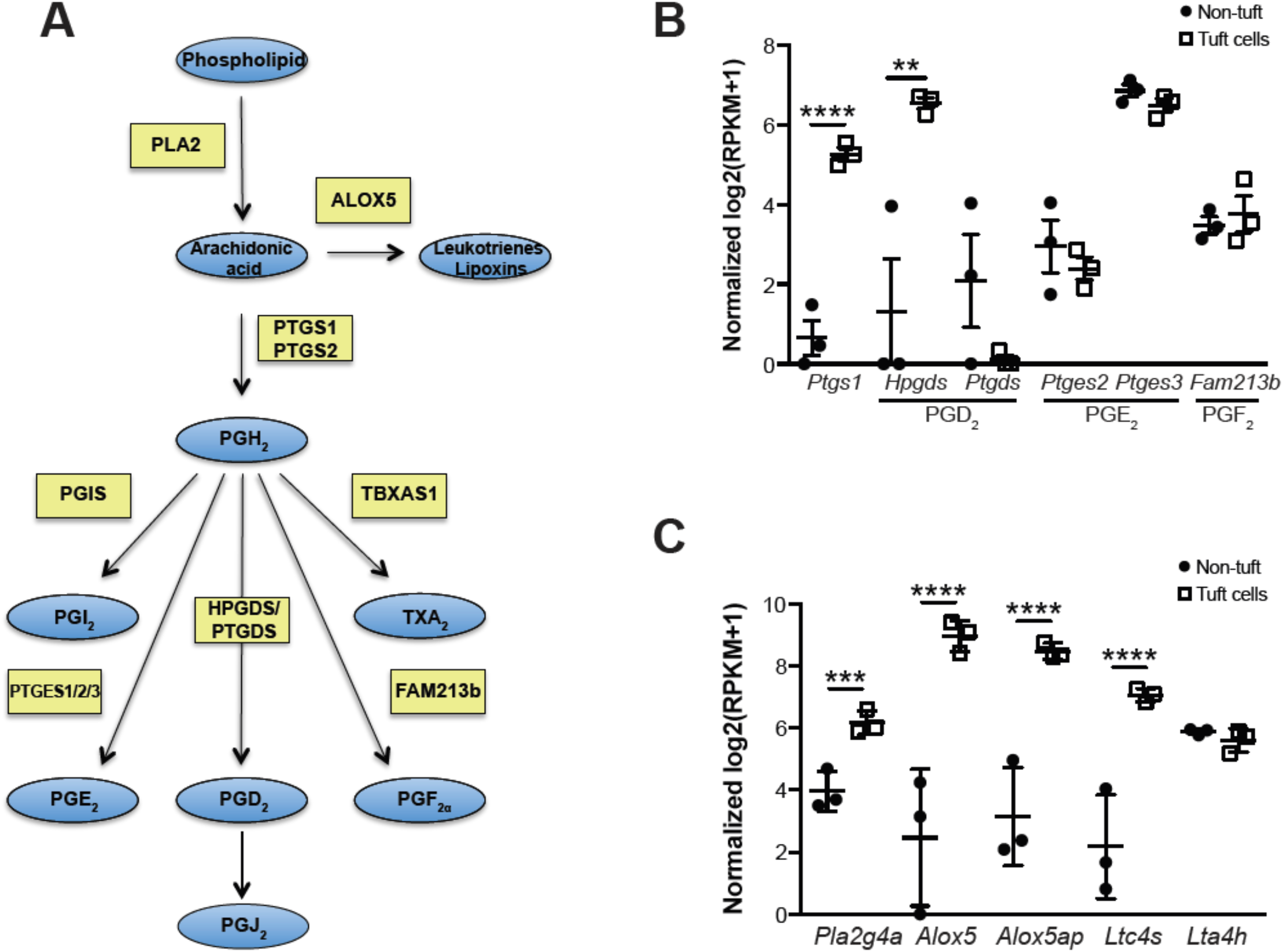
*KC* tuft cells express prostaglandin and lipoxin/leukotriene synthases. (**A**) Diagram of the eicosanoid production pathway leading to synthesis of inflammatory mediators leukotrienes, lipoxins, and prostaglandins. Blue ovals, lipids; yellow boxes, enzymes. Modified from Dennis *et al*. (**B**) RNA-seq demonstrates significantly higher expression of prostaglandin PGD_2_ synthase, Hpgds, and (**C**) the entire pathway of enzymes required to produce leukotriene LTC_4_ in *KC* tuft cells. n = 3. **, p<0.01; ***, p<0.005; ****, p<0.001.

**Figure S11.**
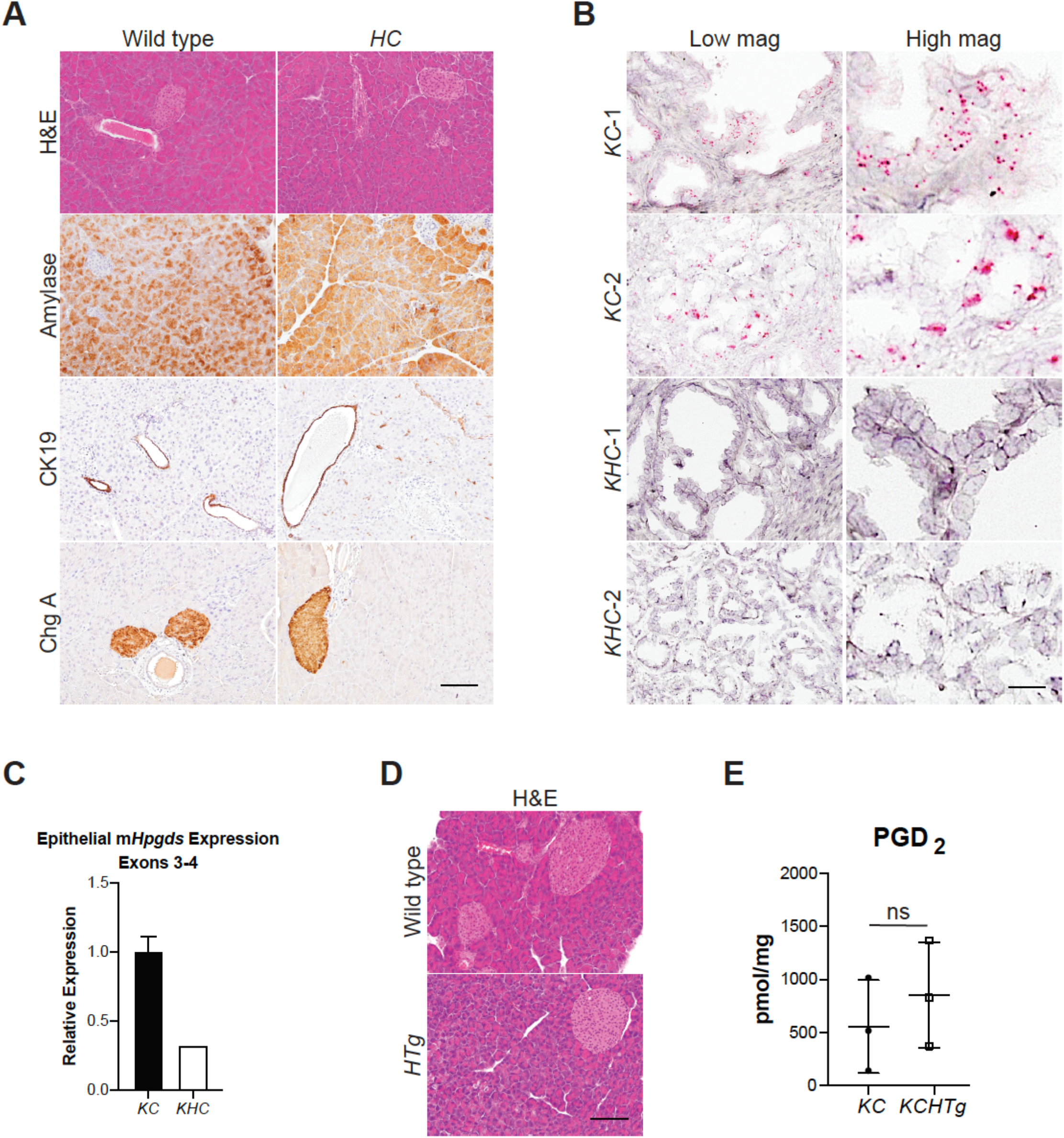
Hpgds ablation or over-expression does not affect pancreas architecture or tuft cell formation. (**A**) Histological analyses of wild type and *Hpgds^fl/fl^;Ptf1a*^*Cre*/+^ (*HC*) mice. Amylase, acinar cell marker; CK19 (cytokeratin 19), ductal marker; ChgA (chromagranin A), islet marker. Scale bar, 100 μm. (**B**) RNA expression analysis by in situ hybridization of exons 2-3 of murine *Hpgds* (pink). Scale bar, 50μm for low magnification images (low mag); 25μm for high magnification images (high mag). (**C**) RT-qPCR analysis of a region spanning partial exons 3-4 of murine *Hpgds* in the epithelium of *KC* and *KHC* mice. (**D**) H&E analysis of 6-month-old wild type or *Hpgds-Tg* (*HTg*) mice. Scale bar, 100 μm. (**E**) PGD_2_ levels in 6-month-old *KC* or *KCHTg* pancreata. ns, not significant.

**Figure S12.**
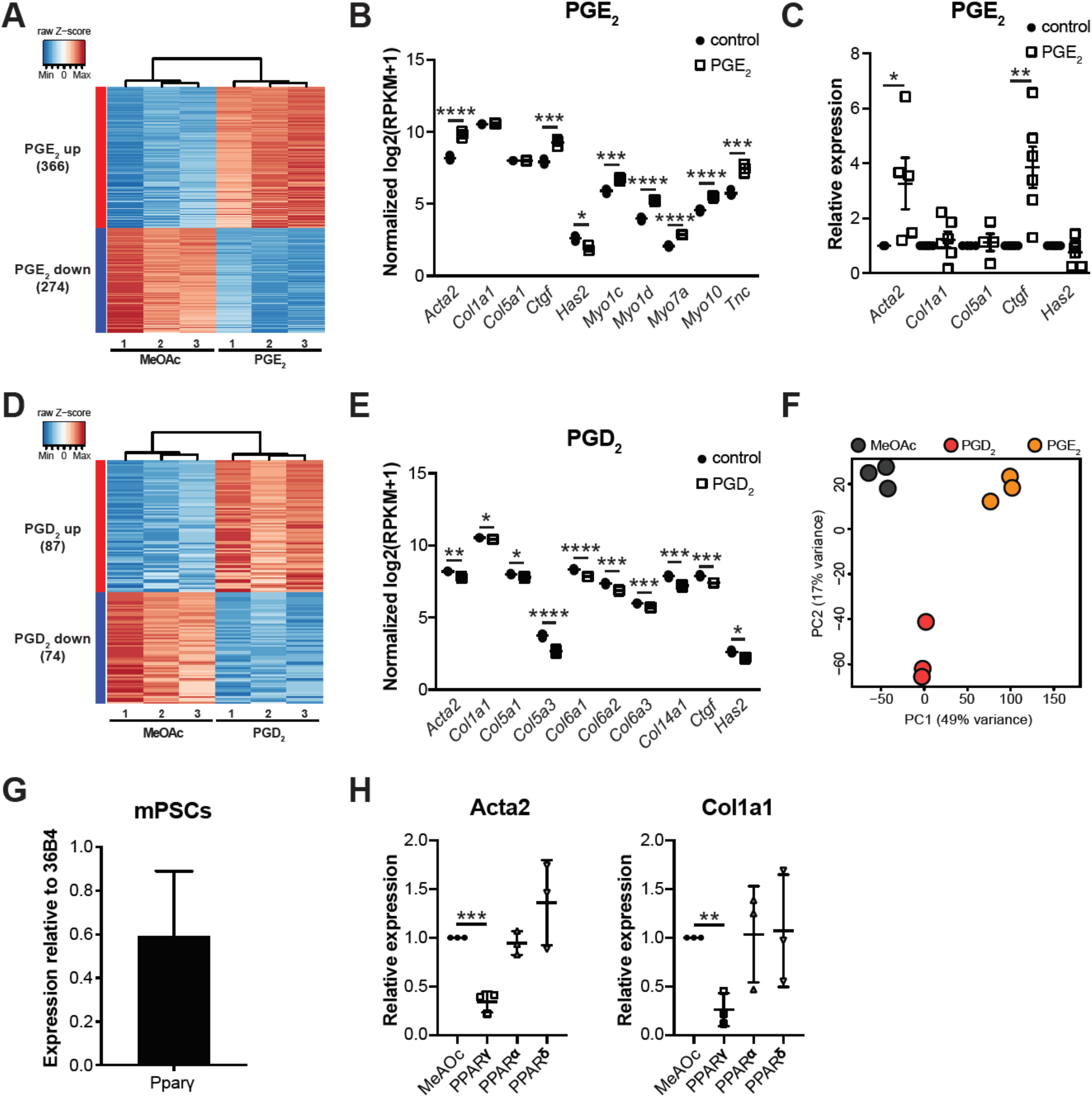
Prostaglandins impact pancreatic stellate cell activation. (**A**) Heat map with hierarchial clustering showing differentially expressed genes in PGE_2_-treated mPSCs vs. control (MeOAc). (**B**) RNA-seq demonstrating an increase in a number of fibrosis-related genes in response to PGE_2_ treatment, confirmed by RT-qPCR in (**C**). (**D**) Heat map with hierarchial clustering showing differentially expressed genes in PGD_2_-treated mPSCs v. control. (**E**) RNA-seq demonstrating a decrease in a number of fibrosis-related genes in response to PGD_2_ treatment. (**F**) Principal Component Analysis comparing the transcriptomes of mPSCs treated with PGD_2_ vs. PGE_2_ vs. control. n = 3 mice per group. (**G**) RT-qPCR for Pparγ in MeAOC-treated mPSCs. (**H**) Pparγ, but not Pparα or Pparδ agonists phenocopy PGD_2_ treatment. *, p<0.05; **, p<0.01; ***, p<0.005; ****, p<0.001.

**Figure S13.**
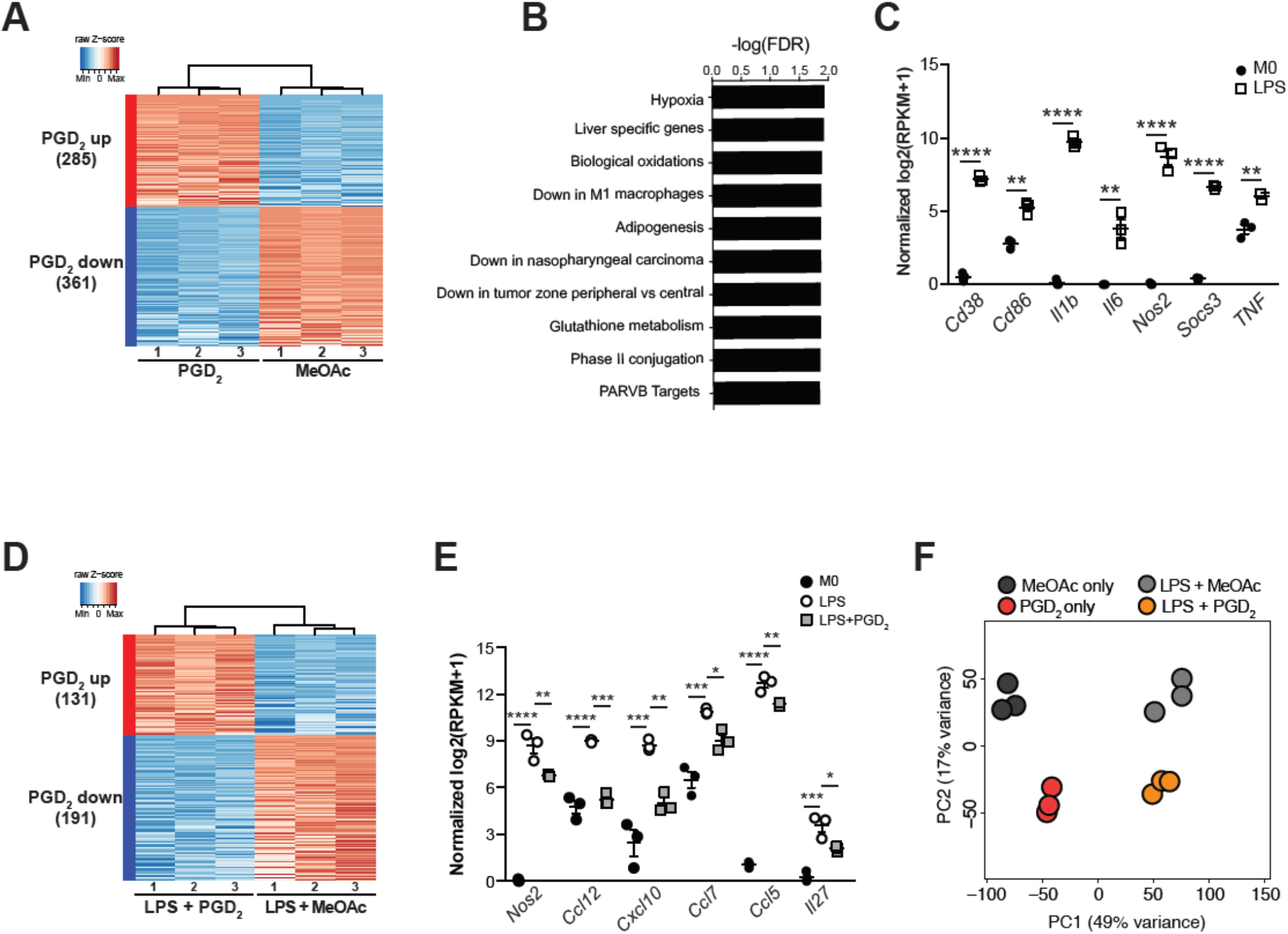
PGD_2_ impacts macrophage polarization. (**A**) Heat map with hierarchical clustering showing differentially expressed genes (p<0.01 and average fold change>1.5) in MeAOc or PGD_2_-treated bone marrow macrophages (BMM). (**B**) GSEA analysis of pathways up-regulated in response to PGD_2_ treatment. (**C**) RNA-seq demonstrates a significant increase in genes associated with pro-inflammatory macrophages in response to LPS treatment. (**D**) Heat map with hierarchical clustering showing differentially expressed genes (p<0.01 and average fold change>1.5) in LPS and MeAOc or PGD_2_-treated BMM. (**E**) RNA-seq shows a significant decrease in expression of several cytokines in LPS-stimulated BMMs treated with PGD_2_. (**F**) Principal Component Analysis comparing the transcriptomes of bone marrow macrophages (BMM) treated with PGD_2_ vs. control or PGD_2_ + LPS vs. MeAOc control. n = 3 mice per group. *, p<0.05; **, p<0.01; ***, p<0.005; ****, p<0.001.

**Figure S14.**
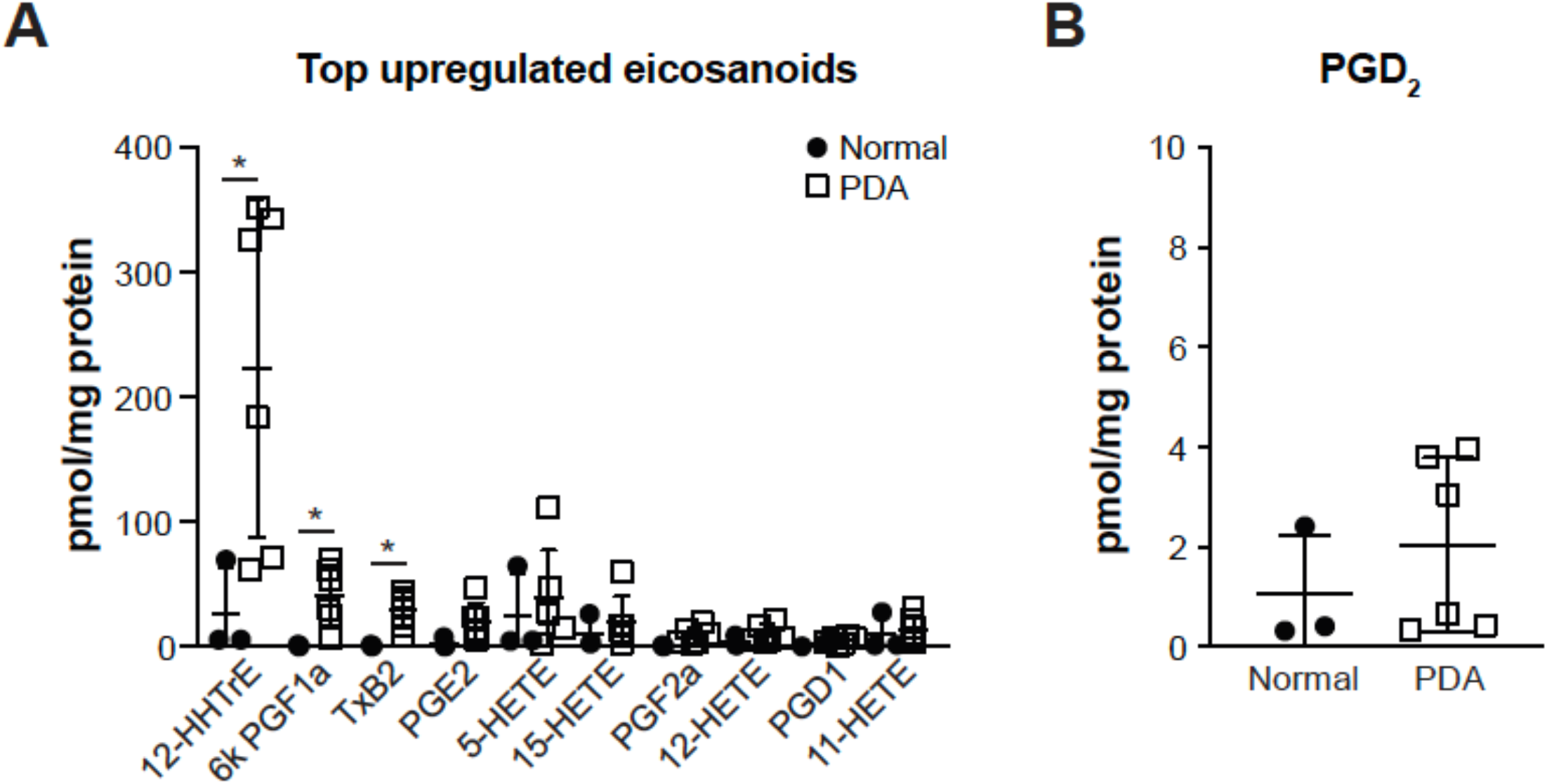
Eicosanoids in human pancreas and pancreatic ductal adenocarcinoma. (**A**) Eicosanoid expression profiling of whole tissue from human pancreatic ductal adenoma (n = 6) vs. nearby normal tissue (n = 3). The top 10 most highly differentially expressed eicosanoids are shown. (**B**) PGD_2_ expression in these same samples. *, p < 0.05.

**Table S1.** Primary antibodies and binding proteins used in immunofluorescence (IF) and immunohistochemistry (IHC) studies.

| Antibody | Company | Catalog # | Dilution – IF | Dilution – IHC |
| --- | --- | --- | --- | --- |
| αSMA | Santa Cruz | sc-32251 | N/A | 1:1000 |
| Amylase | Abcam | ab21156 | N/A | 1:5000 |
| Bodipy | Thermo Fisher | D3922 | 1:100 | N/A |
| Cd3 | Abcam | ab5690 | N/A | 1:1000 |
| Cd45 | Biolegend | 103101 | N/A | 1:100 |
| Ck19 | Epitomics | AC-0073 | N/A | 1:200 |
| Cleaved<br>Caspase 3 | Cell Signaling | 9661S | N/A | 1:1000 |
| Cox1 | Santa Cruz | sc-1754 | 1:100 | 1:500 |
| Dclk1 | Abcam | ab37994 | 1:200 | 1:500 |
| EGFR (Y1092) | Abcam | ab40815 | 1:100 | N/A |
| F4/80 | Biorad | MCA497R | N/A | 1:1000 |
| HABP | Calbiochem | 385911 | N/A | 1:500 |
| Hpgds | Cayman<br>Chemical | 160013 | 1:500 | 1:1000 |
| Ki67 | Abcam | ab15580 | 1:4000 | 1:8000 |
| Pou2f3 | Santa Cruz | Sc-330 | 1:100 | 1:5000 |
| RFP | MBL Life<br>science | PM005 | 1:1000 | N/A |
| Siglec f | BD Biosciences | 552125 | 1:100 |  |
| Trpm5 | Novus<br>Biologicals | NB100-98867 | 1:1000 | 1:20000 |
| Vav1 | Cell Signaling | 2502 | 1:100 | 1:200 |

**Table S2.**
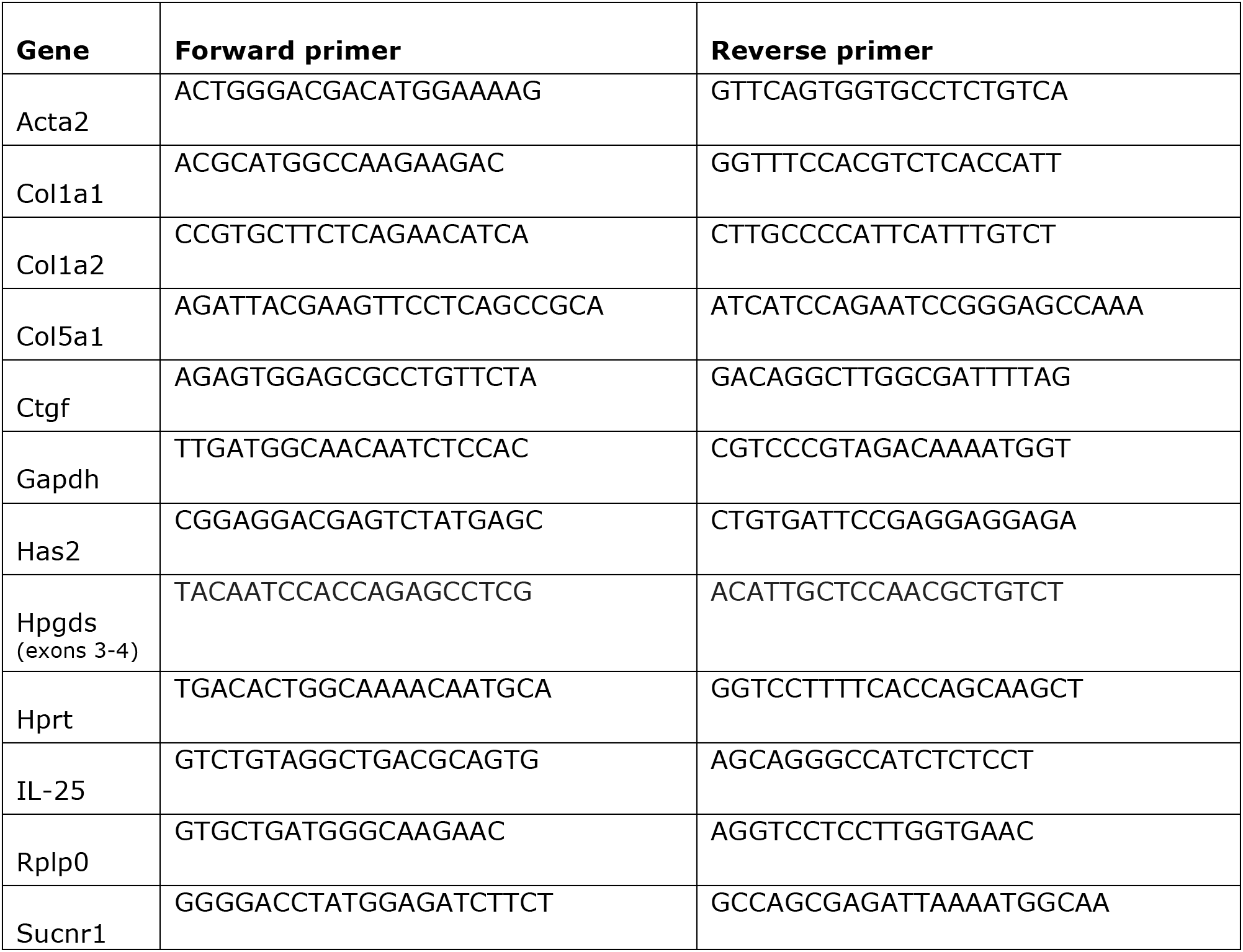
Primers used for RT-qPCR analyses.

**Movie S1. 3-D reconstruction of a PanIN tuft cell** including nucleus-associated, lavender, and cytoplasmic, yellow, lipid droplets, highlighting their organization with the cell. Microvilli, blue; nucleus, red; basement membrane, orange.

**Movie S2. Nuclear association of tuft cell lipid droplets.** A close-up view of tuft cell lipid droplets making contact with the nucleus, lavender. Cytoplasmic lipid droplets, yellow; Microvilli, blue; nucleus, red.

**Supplemental File 1** Tuft cell Smart-seq2 analysis

**Supplemental File 2** Eicosanoid profiling, WT vs. *KC* pancreata

**Supplemental File 3** RNA-seq mPSCs eicosanoids

**Supplemental File 4** RNA-seq BMMs eicosanoids

**Supplemental File 5** Eicosanoid profiling, human PDA and near-by normal pancreas

